# Classifying T cell activity in autofluorescence intensity images with convolutional neural networks

**DOI:** 10.1101/737346

**Authors:** Zijie J. Wang, Alex J. Walsh, Melissa C. Skala, Anthony Gitter

## Abstract

The importance of T cells in immunotherapy has motivated developing technologies to better characterize T cells and improve therapeutic efficacy. One specific objective is assessing antigen-induced T cell activation because only functionally active T cells are capable of killing the desired targets. Autofluorescence imaging can distinguish T cell activity states of individual cells in a non-destructive manner by detecting endogenous changes in metabolic co-enzymes such as NAD(P)H. However, recognizing robust patterns of T cell activity is computationally challenging in the absence of exogenous labels or information-rich autofluorescence lifetime measurements. We demonstrate that advanced machine learning can accurately classify T cell activity from NAD(P)H intensity images and that those image-based signatures transfer across human donors. Using a dataset of 8,260 cropped single-cell images from six donors, we meticulously evaluate multiple machine learning models. These range from traditional models that represent images using summary statistics or extract image features with CellProfiler to deep convolutional neural networks (CNNs) pre-trained on general non-biological images. Adapting pre-trained CNNs for the T cell activity classification task provides substantially better performance than traditional models or a simple CNN trained with the autofluorescence images alone. Visualizing the images with dimension reduction provides intuition into why the CNNs achieve higher accuracy than other approaches. However, we observe that fine-tuning all layers of the pre-trained CNN does not provide a classification performance boost commensurate with the additional computational cost. Our software detailing our image processing and model training pipeline is available as Jupyter notebooks at https://github.com/gitter-lab/t-cell-classification.

## Introduction

Immunotherapy is a type of cancer treatment that uses the body’s own immune cells to boost natural defenses against cancer. T cells are a promising target for immunotherapies because of their antigen specificity and diverse cytotoxic and immune-modulating activities. Prior to activation by an antigen, T cells are in a resting or quiescent state. Upon activation, T cells increase in size, proliferate, and produce cytokines^1^. T cells are highly heterogeneous, due to various states of activation and production of cytokines with cytotoxic or immune-modulating effects. Immunotherapies that enhance T cell cytotoxicity are currently used in clinical cancer treatments^2^. Other immunotherapies that enhance T cell regulatory activities are in development for diseases including HIV and diabetes^3,4^.

Extensive functional heterogeneity at the single-cell level has been observed *in vivo* for CAR T cell immunotherapy for B cell lymphoma in mice, with only approximately 20% of CAR T cell-B cell interactions leading to target killing^5^. The difference in killing efficiency is likely due to heterogeneity in cytotoxic potential among the CAR T cells^5^. Due to this T cell heterogeneity, T cell activation and function must be assessed at the single-cell level to fully characterize and optimize immunotherapy effects on T cells. However, most current T cell profiling methods, such as immunofluorescence of surface protein expression and cytokine production, rely on exogenous contrast agents. Labelling intracellular cytokine production requires cell fixation, limiting application of these methods to *in vitro* assessment of subsets of T cells. A label-free and non-destructive pipeline to determine T cell activation is necessary for *in vitro* characterization and sorting of expanded T cells to ensure optimally functional T cells are used in immunotherapies and for *in vivo* pre-clinical assessment of immunotherapies^6^.

Autofluorescence imaging is appealing because it only relies on endogenous contrast and is non-destructive. Endogenous fluorophores include the metabolic co-enzymes NAD(P)H and FAD as well as collagen. The fluorescence lifetime is the time that the fluorophore is in the excited state, typically picoseconds to nanoseconds in duration, and is sensitive to the microenvironment of the fluorophore. Activated, functional immune cells require specific metabolic programs to support high levels of proliferation and cytokine production. Therefore, autofluorescence imaging of NAD(P)H and FAD provides endogenous endpoints of cellular metabolism reflective of immune cell function. Previous studies have used autofluorescence lifetime imaging to identify macrophages within the tumor microenvironment *in vivo*^7^ and classified activation state of T cells *in vitro*^6^. The fluorescence lifetime of NAD(P)H and FAD is highly sensitive to the microenvironment and binding of NAD(P)H and FAD. Thus, this fluorescence lifetime can be used to resolve metabolic differences between functional states of immune cells. However, fluorescence lifetime imaging requires specialized and expensive microscope components, limiting its use to particular labs. Although autofluorescence intensity images lack the depth of information provided by the fluorescence lifetime, intensity images can easily and quickly be acquired on almost any commercial fluorescence microscope, allowing widespread adoption and seamless integration of a new technique into existing protocols of live cell assessment. Here, we develop a computational framework that uses autofluorescence intensity images to assess T cell activation state at the single-cell level.

Machine learning is promising for classifying cell subtypes from label-free images. For example, Pavillon et al. used regularized logistic regression to predict macrophage activation state^8^, and Yoon et al. identified lymphocyte cell types with k-nearest neighbors^9^. Advanced machine learning models, in particular convolutional neural networks (CNNs), are now the prevailing approach for a variety of cellular image analyses^10–12^. CNNs can classify cell phenotypes^13,14^, segment cells^15^, and predict protein localization^16, 17^, cell lineage choice^18^, or ratios of activated T cells in a population^19^. They are also effective at cell type classification tasks such as differentiating T cells from cancer cells^20^, predicting cell cycle state^21^, and cell sorting^22^.

In this study, we use transfer learning with a pre-trained CNN to classify T cell activation state at the single-cell level. Transfer learning re-uses a model for one task to improve performance on another task. Instead of extracting a small set of features from images before training a cell type classifier^20^, we treat the autofluorescence intensity images as the input and take advantage of an existing CNN that has been trained on generic images. The pre-trained CNN extracts high dimensional image features. We train a simple classifier on these features or fine-tune partial layers to adapt the CNN for T cell activity classification. Repurposing a CNN pre-trained on generic images has been successful in medical imaging applications^23–25^ and cellular image analyses^26^ such as classifying white blood cell types^27^, recognizing cell staining patterns^28^, and predicting mechanism of action in compound treatments^29–31^. Compared to end-to-end CNN training, the transfer learning approach is more computationally efficient and requires fewer training samples. Because T cells differ from donor to donor in real-life immunotherapy applications, we use a rigorous donor-specific cross-validation scheme to train and evaluate our models. For the same reason, we hold out all images from one donor and only use them to assess the final performance of our best model.

The pre-trained CNN can accurately classify T cell activity across donors with autofluorescence intensity images as the only input. We compare the pre-trained CNN to a spectrum of simpler models to better understand when and why deep learning is needed. Adapting pre-trained CNNs is an important strategy in this domain and the most accurate approach overall, improving upon classifiers that operate on pre-extracted cell image features. In particular, fine-tuning some higher-level layers outperforms directly using pre-trained CNN extracted features. However, it is generally not worth the additional computational expense to fine-tune all layers of the CNN. Interpretation techniques demonstrate that the pre-trained CNN learns better representations for the two types of T cell images than other classifiers. Our success in classifying T cell activity without exogenous contrast agents or fluorescence lifetime suggests that modern machine learning approaches may help compensate for imaging data with less molecular specificity.

## Results

### Overview

Our goal is to classify individual T cells as activated (positive instances) or quiescent (negative instances) using only cropped autofluorescence intensity cell images. We explore multiple classification approaches of increasing complexity. A frequency classifier uses the frequency of positive samples in the training set as the probability of the activated label. This naive baseline model assesses how well the class skew in the training images can predict the label of new images. In addition, we test three Lasso logistic regression approaches on different featurizations of the cropped T cell images. The first uses the image pixel intensities directly as features. The second uses only two image summaries as features, the cell size and total intensity. The third uses attributes calculated with CellProfiler^32^, such as the mean intensity value and cell perimeter.

We also assess multiple types of neural networks. A fully connected neural network (multilayer perceptron) generalizes the logistic regression model with pixel intensities by adding a single hidden layer. The LeNet CNN architecture^33^ learns convolutional filters that take advantage of the image structure of the input data. This CNN is simple enough to train from random initialization with a limited number of images. Finally, we consider two deeper and more complex CNNs. Both use transfer learning to initialize the Inception v3 CNN architecture with a model that has been pre-trained on generic (non-biological) images. One version trains a new fully connected layer from scratch using off-the-shelf features extracted from cell images with the pre-trained CNN. An alternative fine-tunes multiple layers of the pre-trained CNN.

The overall workflow for our pre-trained CNN model is described in Fig. 1. The original microscopy images are segmented, cropped, and padded. We filter images that do not contain a T cell and other artifacts, leaving the final image counts for each of the six donors shown in Table 1. Then we train, evaluate, and interpret the machine learning models. Fig. 1 shows the training procedure for the pre-trained CNN as an example.

**Table 1.** Image count per donor before and after filtering cells. This filter discards images that fail to pass entropy and total intensity thresholds.

|  | Donor 1 | Donor 2 | Donor 3 | Donor 4 | Donor 5 | Donor 6 |
| --- | --- | --- | --- | --- | --- | --- |
| Activated cell count before filtering | 609 | 1139 | 604 | 789 | 791 | 531 |
| Quiescent cell count before filtering | 2184 | 399 | 2351 | 2110 | 528 | 1007 |
| Activated cell count after filtering | 235 | 647 | 446 | 482 | 683 | 442 |
| Quiescent cell count after filtering | 1551 | 141 | 1238 | 1569 | 246 | 580 |

**Figure 1.**
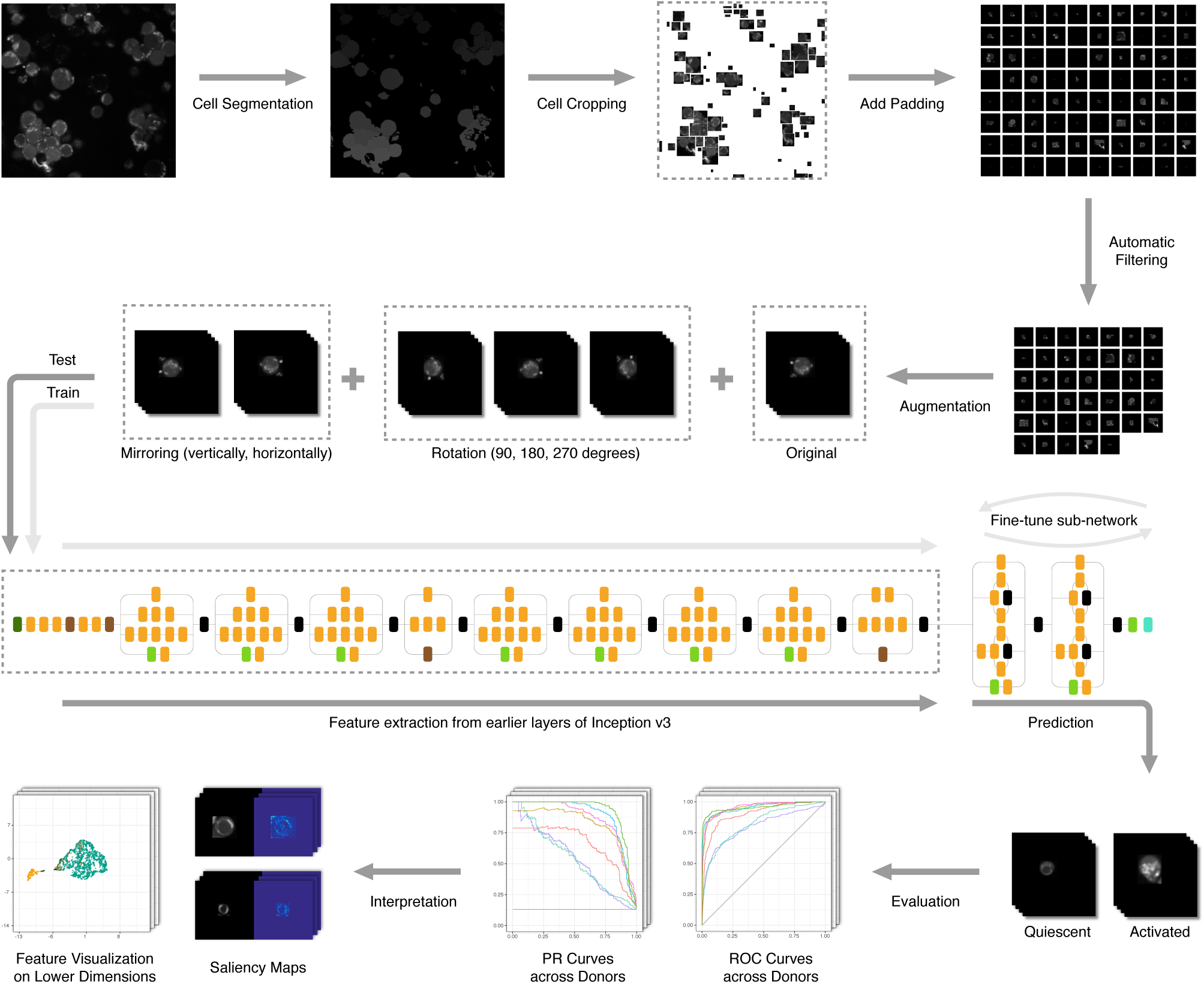
Our T cell image data processing workflow.

The T cell microscopy images may vary from donor to donor. A trained model must be able to generalize to new donors in order to be useful in a practical pre-clinical or clinical setting. Therefore, all of our evaluation strategies train on images from some donors and evaluate the trained models on separate images from a different donor, which is referred to as subject-wise cross-validation^34^ or a leave-one-patient-out scheme^28^. We initially assess the classifiers with cross-validation across donors. In addition, we hold out all images from a randomly selected donor, donor 4, and only use them after completing the rest of our study to confirm that our model selection and hyper-parameter tuning strategies generalize to a new donor.

### Cross-validation Across Donors

In order to assess our classifiers’ performance on cell images from new donors, we design a nested cross-validation scheme to train, tune, and test all models. Due to this cross-validation design, the same model could have different optimal hyper-parameters for different test donors. Therefore, we group the final model performance by test donors (Fig. 2). We plot multiple evaluation metrics because each metric rewards different behaviors^35^. The area under the curve (AUC) and average precision are summary statistics of the receiver operating characteristic (ROC) curve (Fig. 3) and precision recall (PR) curve (Fig. 4), respectively. For all three evaluation metrics, the two pre-trained CNN models outperform other classifiers.

**Figure 2.**
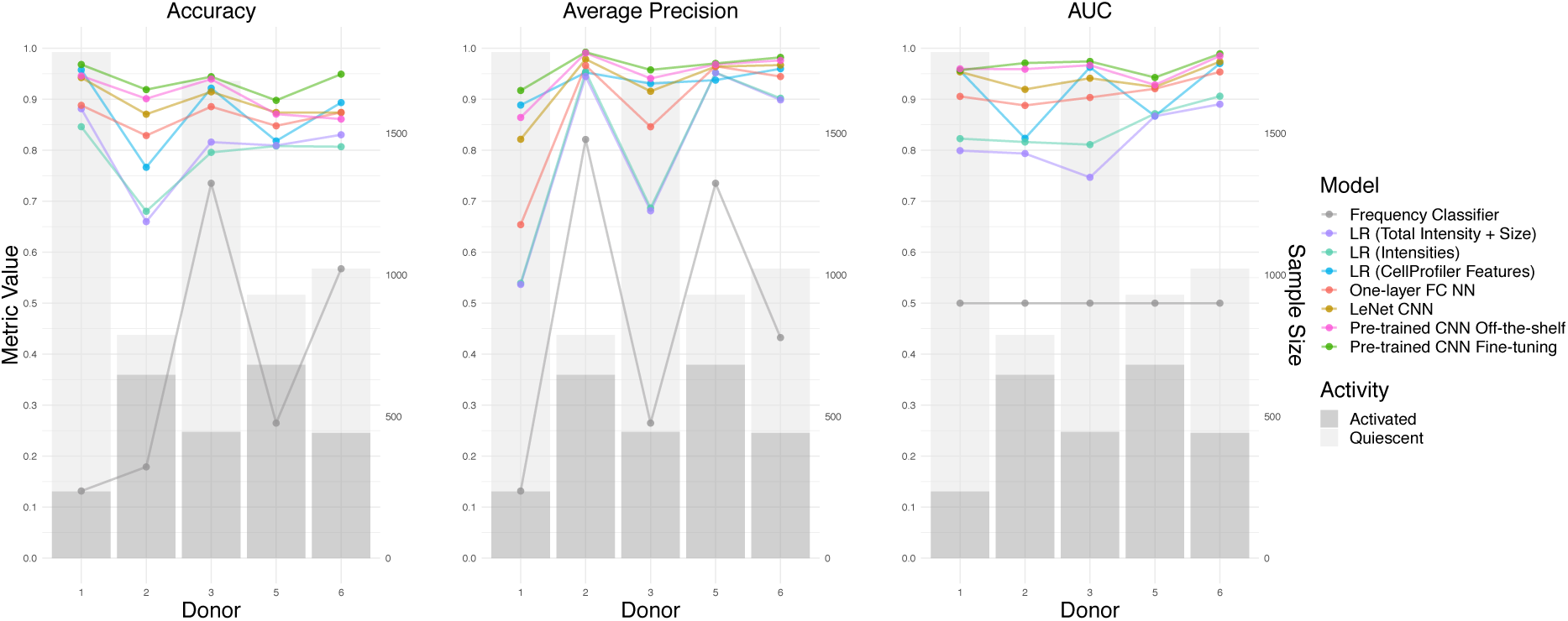
Model summary per donor for three different evaluation metrics. The line graphs display the classifiers’ performance across donors. The bar graphs display the number of activated and quiescent images for each donor, which affects the baseline accuracy and average precision of a random classifier.

**Figure 3.**
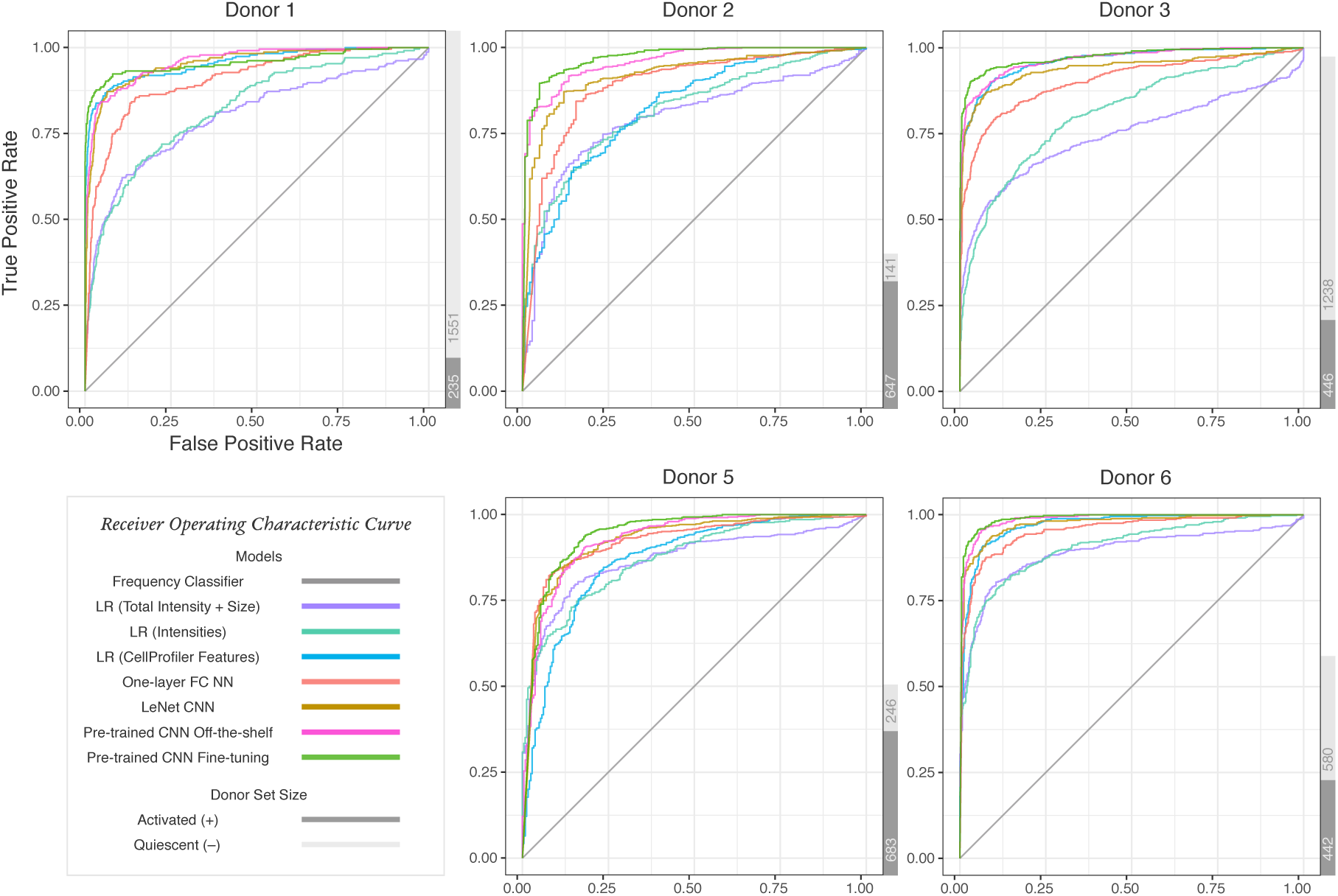
ROC curves for each type of classifier and donor. The gray bars to the right display the number of activated and quiescent images for each donor.

**Figure 4.**
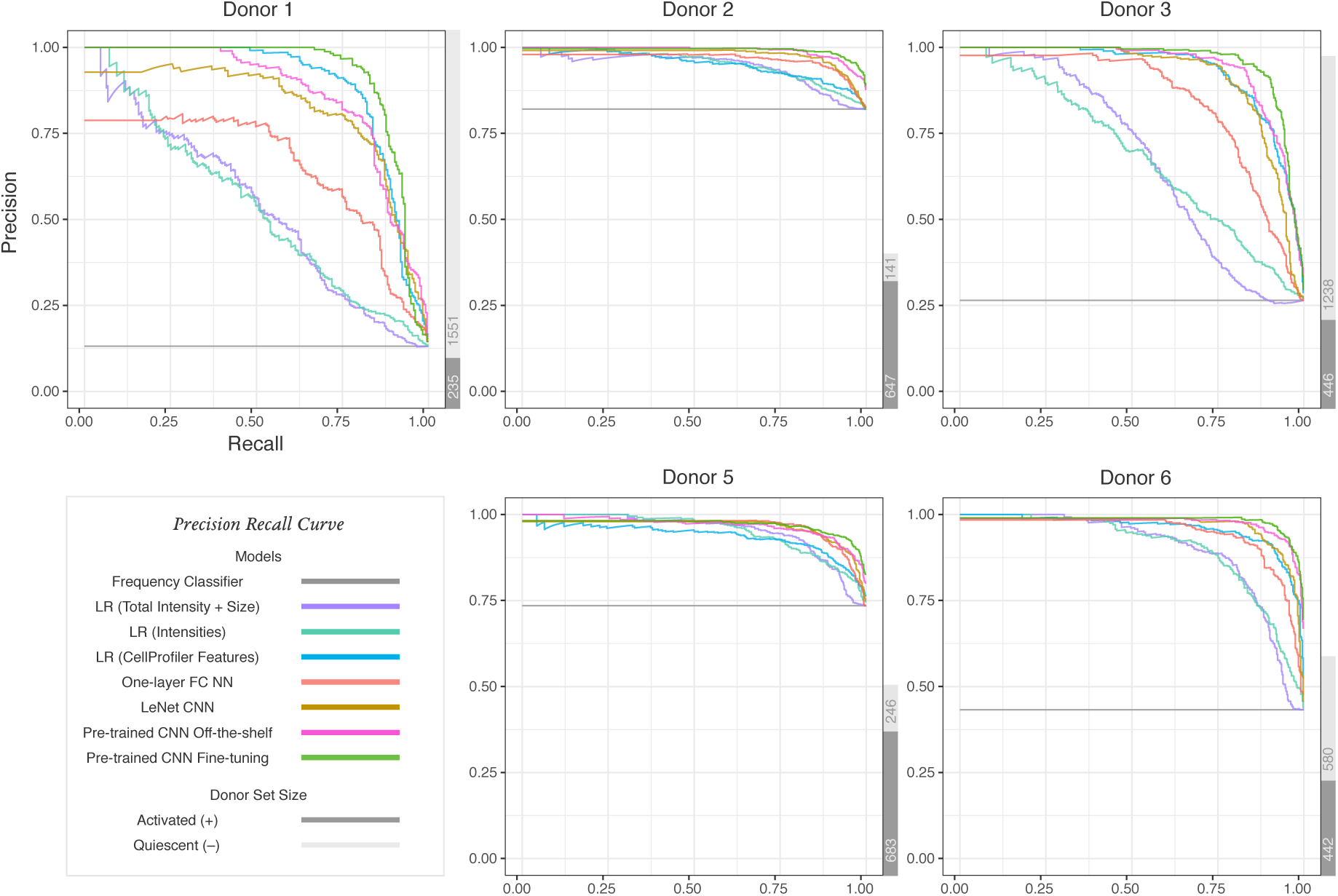
PR curves for each type of classifier and donor. The gray bars to the right display the number of activated and quiescent images for each donor.

The frequency classifier’s average accuracy for all test donors is 37.56% (Fig. 2 and Table S1). The low accuracy of this simple method implies that the majority class in the training and test sets is likely to be different. For example, there are more activated cells from donor 1 while there are more quiescent cells from the combination of donors 2, 3, 5, and 6. This baseline establishes that classifiers that fail to use features other than the label count will perform poorly.

Three logistic regression models using different features all give better classifications than the baseline model. Logistic regression with the image pixel matrix leads to an average accuracy of 78.74% (Fig. 2 and Table S2). Among those 6724 pixel features, 5822 features on average are removed by the Lasso regularization. To interpret this model, we plot the exponential of each pixel’s coefficient to visualize the odds ratios. As shown in Fig. S1, this model learns the shape of cells. Larger cells are more likely to be classified as activated. Logistic regression using only mask size and total intensity as features gives slightly better performance with an average accuracy of 79.93% (Fig. 2 and Table S3). For all test donors, the optimal coefficient of cell mask size is negative, whereas the coefficient of total intensity is positive. In practice, we expect larger cells to be activated, but the negative coefficient indicates the model learns the wrong relationship of cell size and activity state. This can be explained by the inconsistent cell size distribution across donors (Fig. S2) and the correlation of cell size and total intensity (multicollinearity). Comparing the odds ratio of one standard deviation increment of each feature, however, shows this logistic regression model is much more sensitive to total intensity than cell size. Finally, the logistic regression model with CellProfiler attributes yields 87.14% average accuracy (Fig. 2 and Table S4). After computing the odds ratio adjusting to the standard deviation of each feature, attributes that are related to image intensity and cell area have the strongest impact on the predictions.

Non-linear models with image pixels as input have accuracies comparable to the logistic regression model with CellProfiler features. We tune the learning rate, batch size and the number of hidden layer neurons of the simple neural network with one hidden layer. Even though its average accuracy of 86.48% (Fig. 2 and Table S5) is slightly lower than logistic regression with CellProfiler features, it has more stable performance across the test donors. In comparison, the LeNet CNN has a more complex architecture and takes advantage of the image structure of the input data. After selecting the best learning rate and batch size, LeNet reaches an average accuracy of 89.51% (Fig. 2 and Table S6).

Our most advanced models using the pre-trained CNN outperform all other methods. Both versions of the pre-trained CNN use cell images as input and require a previously trained CNN. For one version, we use the pre-trained CNN as a feature extractor and then train a new hidden layer with off-the-shelf features. Alternatively, we fine-tune multiple higher-level layers of the CNN with T cell images. We include the fine-tuned layers as a hyper-parameter. Specifically, we define *n*, ranging from 1 to 11, as the number of last Inception modules in the pre-trained Inception v3 CNN to fine-tune. For example, if *n* = 1, we only fine-tune the last Inception module, whereas we fine-tune all the layers of the Inception v3 CNN when *n* = 11. After tuning *n* along with the other hyper-parameters, we compare the CNN with fine-tuning to the CNN off-the-shelf model in order to study the effect of fine-tuning on classifier performance. Additionally, we compare the test results of different *n* to analyze how the number of fine-tuned layers affects classification.

The average accuracy for the pre-trained CNN off-the-shelf model is 90.36% (Fig. 2 and Table S7) and 93.56% for the pre-trained CNN with fine-tuning (Fig. 2 and Table S8). The fine-tuning model uses 11, 10, 7, 11, and 8 layers as the optimal *n* for the five test donors. However, depending on the test donor and the evaluation metric, the number of fine-tuned layers does not necessarily have a strong effect on the predictive performance (Fig. 5). Different *n* values yield similar evaluation metrics. Fine-tuning all 11 layers also greatly increases the CNN training time (Figs. S3 and S4).

**Figure 5.**
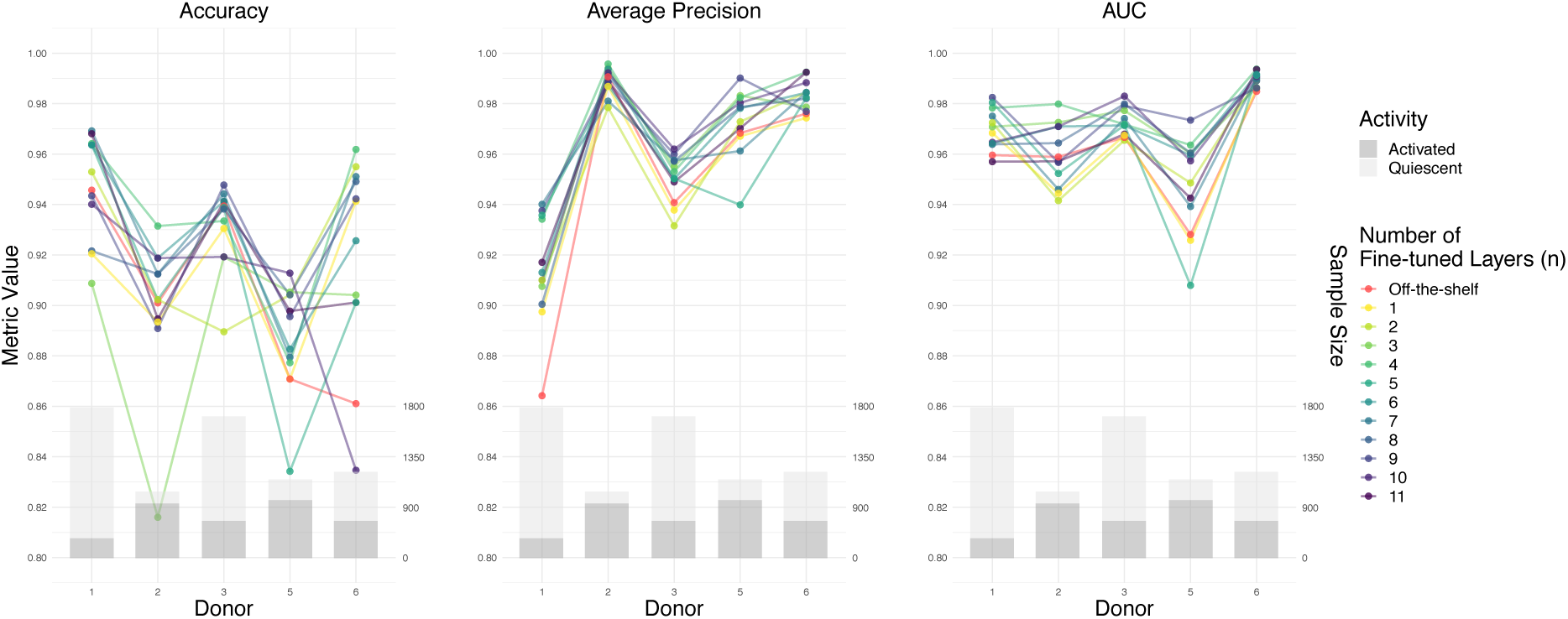
Performance comparison of fine-tuning a different number of layers and the pre-trained CNN off-the-shelf model.

### Confirming Generalization with a New Donor

In order to evaluate our ability to generalize to T cell images from a new individual, we completely hold out images from donor 4 during the study design, model implementation, and cross-validation above. We apply the same nested cross-validation scheme to train, tune, and test the pre-trained CNN with fine-tuning, the most accurate model in the previous cross-validation, on images from donor 4. It gives an accuracy of 98.83% (Table 2). Out of 2,051 predictions, there are only 4 false positives and 20 false negatives. The performance metrics in Table 2 are substantially higher than their counterparts in Table S8. Having training data from five donors instead of four likely contributes to the improved performance.

**Table 2.** Performance of the pre-trained CNN with fine-tuning on held out donor 4.

| Donor | Accuracy | Precision | Recall | Average Precision | AUC | Activated Count | Quiescent Count |
| --- | --- | --- | --- | --- | --- | --- | --- |
| 4 | 98.83% | 99.14% | 95.85% | 99.79% | 99.93% | 482 | 1569 |

### Pre-trained CNN with Fine-tuning Errors

We inspect the T cell images that the pre-trained CNN with fine-tuning classifies incorrectly in order to better understand its failures and accuracy. We visualize the misclassified images for all test donors in Figs. S5-S10 along with the predicted label, the Softmax score of the network output layer, and the temperature scaled confidence calibration^36^. The majority of misclassified cell images are badly cropped, with no cells or multiple cells included in the frame. Therefore, using a more progressive dim image filter or adding a multiple-cell detector in the image processing pipeline could further improve the model performance. However, for other images with a clear single cell in the frame, the pre-trained CNN tends to give high confidence in its misclassification. These scores suggest that these errors cannot be easily fixed without a more powerful classifier or more diverse training dataset. Temperature scaling could either soften the original Softmax score toward 50% or increase the confidence toward 100%. For the misclassified images in our study, temperature scaling always drops the Softmax probability. This observation matches Guo et al.’s finding that neural networks with higher model capacity are more likely to be overconfident in their predictions^36^.

### Pre-trained CNN with Fine-tuning Interpretation

Visualizing the T cell dataset in 2D helps us understand why some classifiers perform better than others. We use Uniform Manifold Approximation and Projection (UMAP)^37^ to project the images into 2D such that similar images in the original feature space are nearby in the 2D space. Coloring the images with their activity labels shows how different input representations or learned representations separate the activated and quiescent cells. For example, in Fig. 6, each dot corresponds to one image based on its representation in the last layer of the pre-trained CNNs with fine-tuning. UMAP projects the 2048 learned features in the last layer of the CNN into 2D. In general, the activated and quiescent cells are well-separated in the 2D space, suggesting that the CNN has successfully learned distinct representations for the two types of T cells. Using t-Distributed Stochastic Neighbor Embedding (t-SNE)^38^ instead of UMAP for dimension reduction provides qualitatively similar results (Fig. S11).

**Figure 6.**
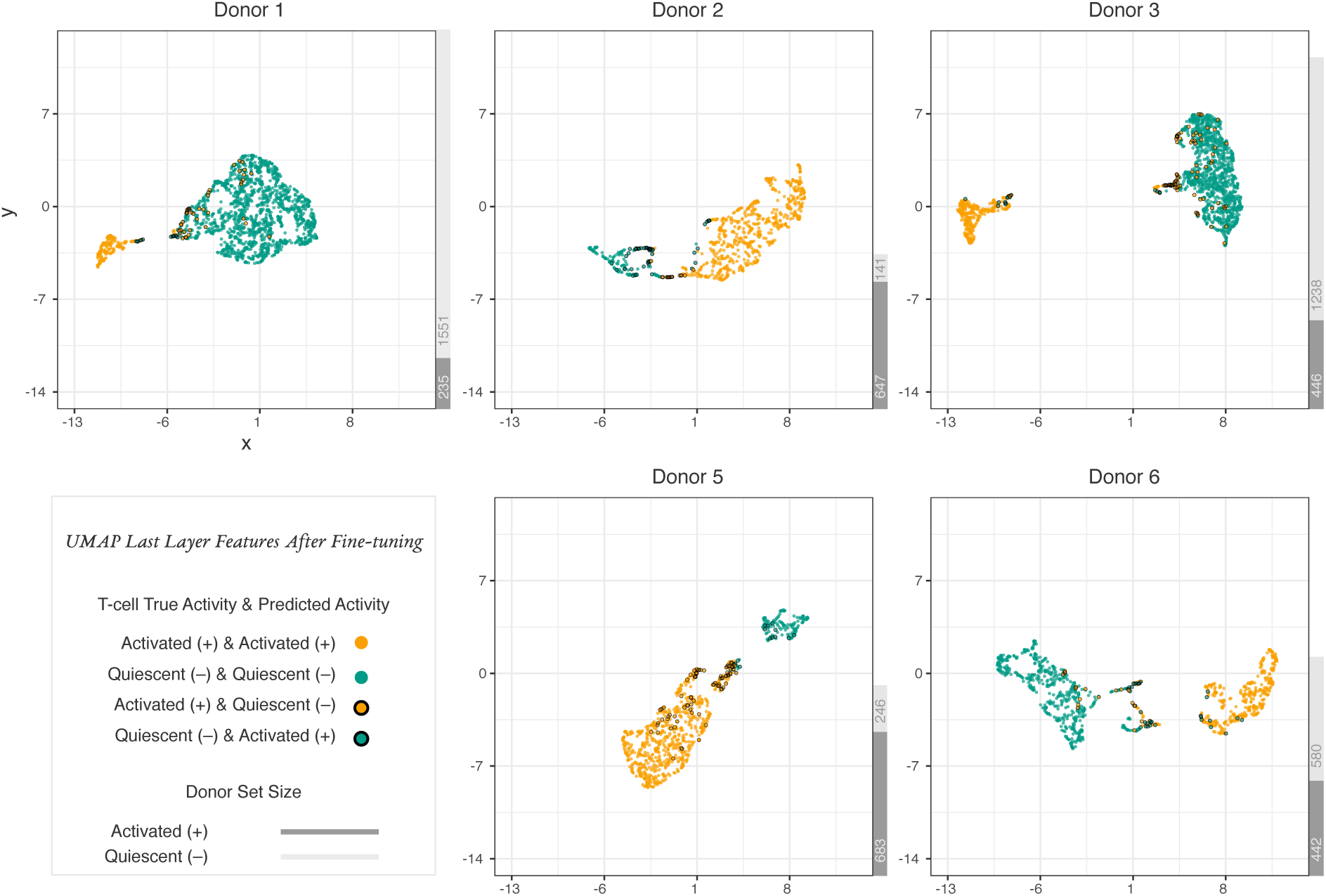
2D representations of T cell features extracted from the pre-trained CNN with fine-tuning. Dimensions are reduced from 2048 using UMAP. The thick outlines indicate incorrect cell activity state predictions.

Generating similar UMAP plots for three alternative image representations shows that the two image classes are not as well separated (Figs. S12-S14). When using the raw pixel features (Fig. S12), the two types of T cells are spread throughout the 2D space. This contributes to the lower performance of the logistic regression and fully connected neural network models that operate directly on pixel intensity. Similarly, there is only moderate spatial separation when using the CellProfiler features (Fig. S13) or the last layer of the CNN before fine-tuning it to predict T cell activity (Fig. S14). These comparisons demonstrate the strong effect of fine-tuning the pre-trained CNN and also help explain the superior performance of pre-trained CNNs over the logistic regression model with CellProfiler features. In addition, by labeling misclassified images as outlined dots in Fig. 6, we see where the pre-trained CNN with fine-tuning makes errors. The incorrect predictions are predominantly distributed in the boundary between the two clusters.

In addition to visualizing the feature representation in the pre-trained CNNs with fine-tuning, we use saliency maps^39^ to interpret how these models make decisions. We generate saliency maps by computing the gradient of the CNN class score with respect to a few randomly chosen donor 1 images from both the activated and quiescent classes (Fig. 7). We use two methods to calculate gradients: standard backpropagation and guided backpropagation^40^. In these heat maps, larger values (green or yellow) highlight the image regions that cause the most change in the T cell activity prediction. Smaller values (dark blue or purple) indicate pixels that have less influence. The uniformly dark blue background in both types of saliency maps indicates that the pre-trained CNNs with fine-tuning have learned to focus on the original cell image instead of the black padding. The larger values in the saliency maps with guided backpropagation often align with the high-intensity regions of the cell images, which correspond to mitochondria and depict metabolic activity^41^. Although the influential regions of the guided backpropagation-based saliency maps are biologically plausible, this type of saliency map is insensitive to random changes of either the input data or model parameters^42^. The saliency maps generated with standard backpropagation are properly affected by these randomized controls but do not concentrate on the high-intensity regions of the input images.

**Figure 7.**
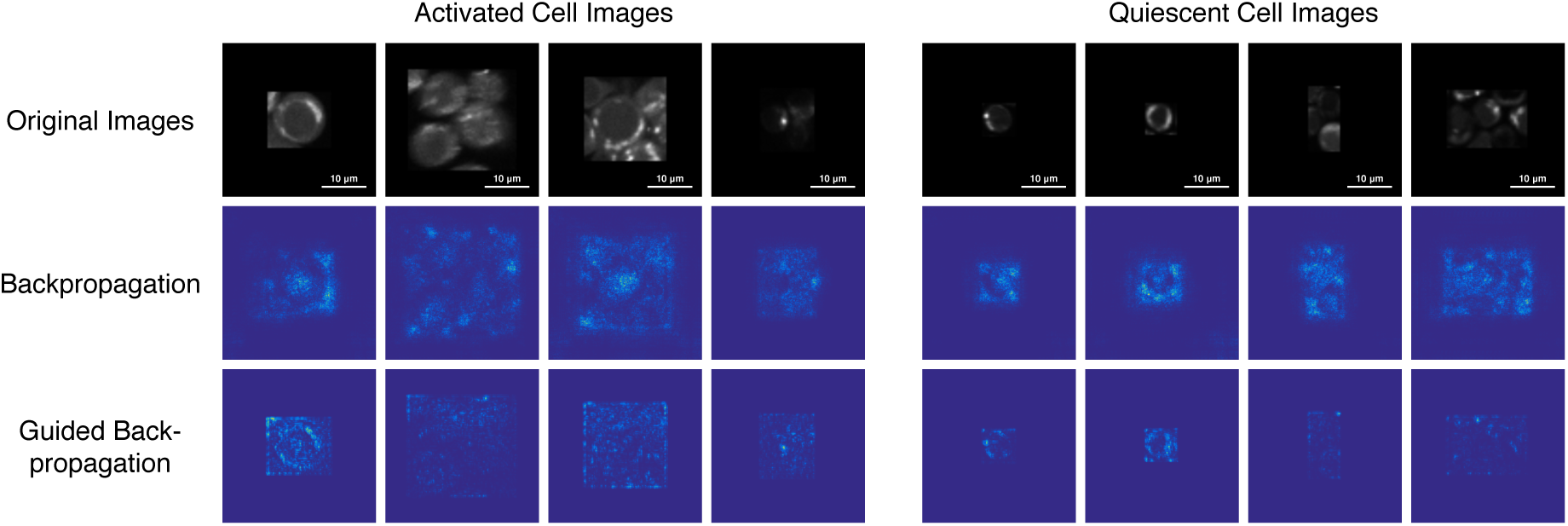
Saliency maps of randomly selected cell images from donor 1 (scale bar: 10 µm). The backpropagation and guided backpropagation rows show two different techniques for generating saliency maps from the same T cell images in the first row.

### Running the Analysis Pipeline

Our GitHub repository https://github.com/gitter-lab/t-cell-classification demonstrates and documents all steps of our analysis pipeline, from pre-processing the cropped images to fine-tuning and interpreting the Inception v3 CNN. Our Python code is presented in Jupyter notebooks^43^, which integrate our code with explanations of its functionality and visualizations of its outputs. These notebooks can serve as a tutorial of best practices in machine learning on autofluorescence microscopy images. We use Binder^44^ to enable readers to execute our Jupyter notebooks in a web browser without having to download any software locally or configure a Python environment. We ensure the Jupyter notebooks continue to run as expected by automatically testing them in Linux, macOS, and Windows with the Travis CI and AppVeyor continuous integration services. Our software re-runs our analyses with a randomly selected 10% of the images. This allows users to quickly train the machine learning models and understand our pipeline.

## Discussion

Our study demonstrates that machine learning models trained on autofluorescence intensity images can accurately classify activated and quiescent T cells across donors. Because autofluorescence images are easier to acquire with standard commercial microscopes compared to fluorescence lifetime images, this workflow has the potential to become a widely applicable approach for live T cell profiling. Fine-tuning a pre-trained CNN is the most powerful classification approach, outperforming alternative machine learning models that are commonly used for microscopy image classification over multiple evaluation metrics. In particular, this CNN applied directly to cropped images has better performance than logistic regression with domain-relevant features extracted by CellProfiler.

We thoroughly explored the effect of fine-tuning more layers of the pre-trained CNN and compared it with the off-the-shelf CNN model. The common transfer learning approach fixes the CNN parameters of the initial network layers, which extract learned features from the images, and trains a simple classifier from scratch that predicts the domain-specific image labels. Our results indicate that fine-tuning pre-trained CNN layers yields better performance than directly using off-the-shelf features. In addition, although fine-tuning more layers tends to give better predictive performance (Fig. 5), it is generally not worth the additional computational time and expense to fine-tune all 11 layers (Figs. S3 and S4). Possible reasons include the limited sample size and relatively homogeneous cell image representations. Given the extra computational costs and implementation challenges, we recommend fine-tuning only the last few layers of a pre-trained CNN for similar autofluorescence microscopy applications. In settings that do require fine-tuning additional layers because the images are more heterogeneous, we suggest taking a larger step size in the layer number hyper-parameter optimization.

The machine learning models recognize image attributes that recapitulate biological domain knowledge. Activated T cells are larger in size^6, 45^. In addition, there are metabolic differences between quiescent and activated T cells^1^, which are evident in the NAD(P)H images. The high intensity regions in the images likely correspond to mitochondria, where the majority of metabolism occurs. It is straightforward to inspect the trained logistic regression model that takes total image intensity and mask size as inputs and observe that it correctly recognizes the relationship between NAD(P)H intensity and activation state.

The parameters of the pre-trained CNN with fine-tuning are not as directly interpretable as the logistic regression model. An additional challenge is that different interpretation techniques provide distinct views of the fine-tuned CNN. Nevertheless, there are some indications in the saliency maps that this CNN also reflects T cell biology. Saliency maps help locate which regions of the input image influence the classification the most. With guided backpropagation, the high-intensity regions of the T cell images tend to be the focal points in the saliency maps. This suggests that the CNN may be sensitive to metabolic differences between quiescent and activated cells and not only changes in cell size. However, guided backpropagation and other more advanced saliency maps were found to be independent of the data, model, and model parameters^42^. The standard backpropagation gradient map is sensitive to these controls, but it focuses more on general cell morphology than the metabolic activity within cells.

Each model in our study is only tuned and evaluated once, which limits our ability to assess the statistical significance of the performance differences across models. Substantial computing time and costs are required for nested cross-validation, especially when fine-tuning multiple layers of the pre-trained CNN (Figs. S3 and S4). The fine-tuning jobs took 5,096 hours (212 days) in total to train on GPUs. Therefore, we are unable train each model multiple times to assess the variability in model performance due to random sampling, computer hardware, non-deterministic algorithms, and other factors. Slight differences in performance should not be over-interpreted.

Based on the misclassified images, the performance of the pre-trained CNN model with fine-tuning is limited by the image cropping quality. Some images contain multiple cells. Others do not contain any T cells. Developing a better filter to detect images with artifacts and adopting state-of-the-art segmentation approaches^15, 46, 47^ could further boost classification accuracy.

Although the pre-trained CNN with fine-tuning has strong predictive performance in this study, there are several caveats regarding how these results may translate to other autofluorescence intensity image classification tasks. We have a small set of donors. The nested cross-validation involves only five donors, and the generalization test uses a single held out donor. In addition, all donors are from a narrow, young, and healthy population. They may not adequately represent the general public or cancer patients, and further study is needed to see if the model is still applicable in more challenging populations. Finally, quantitative fluorescence intensity imaging has technical limitations, requiring consistent imaging settings such as illumination power and detector gain. Future work can assess how robust our trained CNN models are to more diverse imaging settings and whether new training strategies are required to adapt to other domains. For instance, to sort T cells in practice, the classifier would need to be coupled to a flow sorter. The flowing nature of the cells may make the resulting images different enough to require a distinct CNN, which is not an obstacle given recent advances in CNNs for imaging flow cytometry^10, 21, 22, 34, 48, 49^. Overall, our strong results demonstrate the feasibility of classifying T cells directly from autofluorescence intensity images, which can guide future work to bring this technology to pre-clinical and clinical applications.

## Methods

### Cell Preparation and Imaging

This study was approved by the Institutional Review Board of the University of Wisconsin-Madison (#2018-0103). Informed consent was obtained from all donors. The NAD(P)H intensity images in this study were created from a subset of the NAD(P)H fluorescence lifetime images acquired in Walsh et al.^6^. CD3 and CD8 T cells were isolated using negative selection methods (RosetteSep, StemCell Technologies) from the peripheral blood of 6 healthy donors (3 male, 3 female, mean age = 26). The T cells were divided into quiescent and activated groups, where the activated group was stimulated with a tetrameric antibody against CD2, CD3, and CD28 (StemCell Technologies). T cell populations were cultured for 48 hours at 37C, 5% CO2, and 99% humidity.

NAD(P)H intensity images were created by integrating the photon counts of fluorescence lifetime decays at each pixel within the fluorescence lifetime images acquired, as described by Walsh et al.^6^. Briefly, images were acquired using an Ultima (Bruker Fluorescence Microscopy) two-photon microscope coupled to an inverted microscope body (TiE, Nikon) with an Insight DS+ (Spectra Physics) as the excitation source. A 100X objective (Nikon Plan Apo Lambda, NA 1.45), lending an approximate field of view of 110 µm, was used in all experiments with the laser tuned to 750 nm for NAD(P)H two-photon excitation and a 440/80 nm bandpass emission filter in front of a GaAsP photomultiplier tube (PMT; H7422, Hamamatsu). Images were acquired for 60 seconds with a laser power at the sample of 3.0-3.2 mW and a pixel dwell time of 4.6 µs. Grayscale microscopy images were labeled with a deidentified donor ID and T cell activity state according to the culture conditions: quiescent for T cells not exposed to the activating antibodies or activated for T cells exposed to the activating antibodies.

### Image Processing

We segmented cell images using CellProfiler^32^. Each cell was cropped according to the bounding box of its segmented mask. Cell short NAD(P)H lifetime was used to filter out other visually indistinguishable cells (e.g., red blood cells) by removing cells with a mean fluorescence lifetime less than 200 ps. To remove very dim images and images containing no cells, we further filtered the segmented images by thresholding the combination of image entropy and total intensity (Fig. S15). The threshold values in Fig. S15 were chosen based on the distribution of entropy and intensity with a Gaussian approximation. This filter was conservative. We manually inspected the removed images to ensure none of them contained T cells.

Because the classifiers that used image pixels as input required uniform size and some required square images, we padded all activated and quiescent cell images with black borders. The padding size of 82 × 82 was chosen based on the largest image in the dataset after removing extremely large outliers. Also, we augmented the dataset by rotating each original image by 90, 180, and 270 degrees and flipping the original image horizontally and vertically, which added 5 extra images for each cell (Fig. 1). We implemented this image processing pipeline using the Python package OpenCV^50^.

### Nested Cross-validation

We trained and evaluated eight classifiers of increasing complexity (Table 3). We used the same leave-one-donor-out test principle to measure the performance of all models. For example, when using donor 1 as the test donor, the frequency classifier counts the positive proportion among all images in the augmented dataset from donor 2, 3, 5, and 6. Then, it uses this frequency to predict the activity for all unaugmented images from donor 1. By testing in this way, the classification result tells us how well each model performs on images from new donors. Donor 4 was not included in this cross-validation because we randomly selected it as a complete hold-out donor. All images from donor 4 were only used after hyper-parameter tuning and model selection as a final independent test to assess the generalizability of our pipeline to a new donor.

**Table 3.** The eight classifiers with their input features and hyper-parameters.

| Model | Description |
| --- | --- |
| Frequency Classifier | Predict class probability using the class frequencies in the training set. |
| Logistic Regression with Pixel Intensity | Regularized logistic regression model with pixel intensity as input. Regularization power $\lambda$ of $\ell_1$ penalty is tuned. |
| Logistic Regression with Total Intensity and Size | Regularized logistic regression model fitted with two numerical values: image total intensity and cell mask size. Regularization power $\lambda$ of $\ell_1$ penalty is tuned. |
| Logistic Regression with CellProfiler Features | Regularized logistic regression model fitted with 123 features extracted from CellProfiler related to intensity, texture, and area. Regularization power $\lambda$ of $\ell_1$ penalty is tuned. |
| One-layer Fully Connected Neural Network | Fully connected one-hidden-layer neural network with pixel intensity as input. Number of neurons, learning rate, and batch size are tuned. |
| LeNet CNN | CNN with the LeNet architecture with pixel intensity as input. Learning rate and batch size are tuned. |
| Pre-trained CNN Off-the-shelf Model | Freeze layers of a pre-trained Inception v3 CNN. Train a final added layer from scratch with extracted off-the-shelf features. Learning rate and batch size are tuned. |
| Pre-trained CNN with Fine-tuning | Fine-tune the last $n$ layers of a pre-trained Inception v3 CNN. The layer number $n$, learning rate, and batch size are tuned. |

Following the leave-one-donor-out test principle^28, 34^, we wanted the selection of the optimal hyper-parameters to be generalizable to new donors as well. Therefore, we applied a nested cross-validation scheme^51, 52^ (Fig. 8). For each test donor, within the inner loop we performed 4-fold cross-validation to measure the average performance of each hyper-parameter combination (grid search). Each fold in the inner loop cross-validation corresponds to one donor’s augmented images. The outer cross-validation loop used the selected hyper-parameters from the inner loop cross-validation to train a new model with the four other donors’ augmented images. We evaluated the trained model on the outer loop test donor. For models requiring early stopping, we constructed an early stopping set by randomly sampling one-fourth of the unaugmented images from the training set and removing their augmented copies. Then, training continued as long as the performance on images in the early stopping set improved. Similarly, we did not include augmented images in the validation set or the test set.

**Figure 8.**
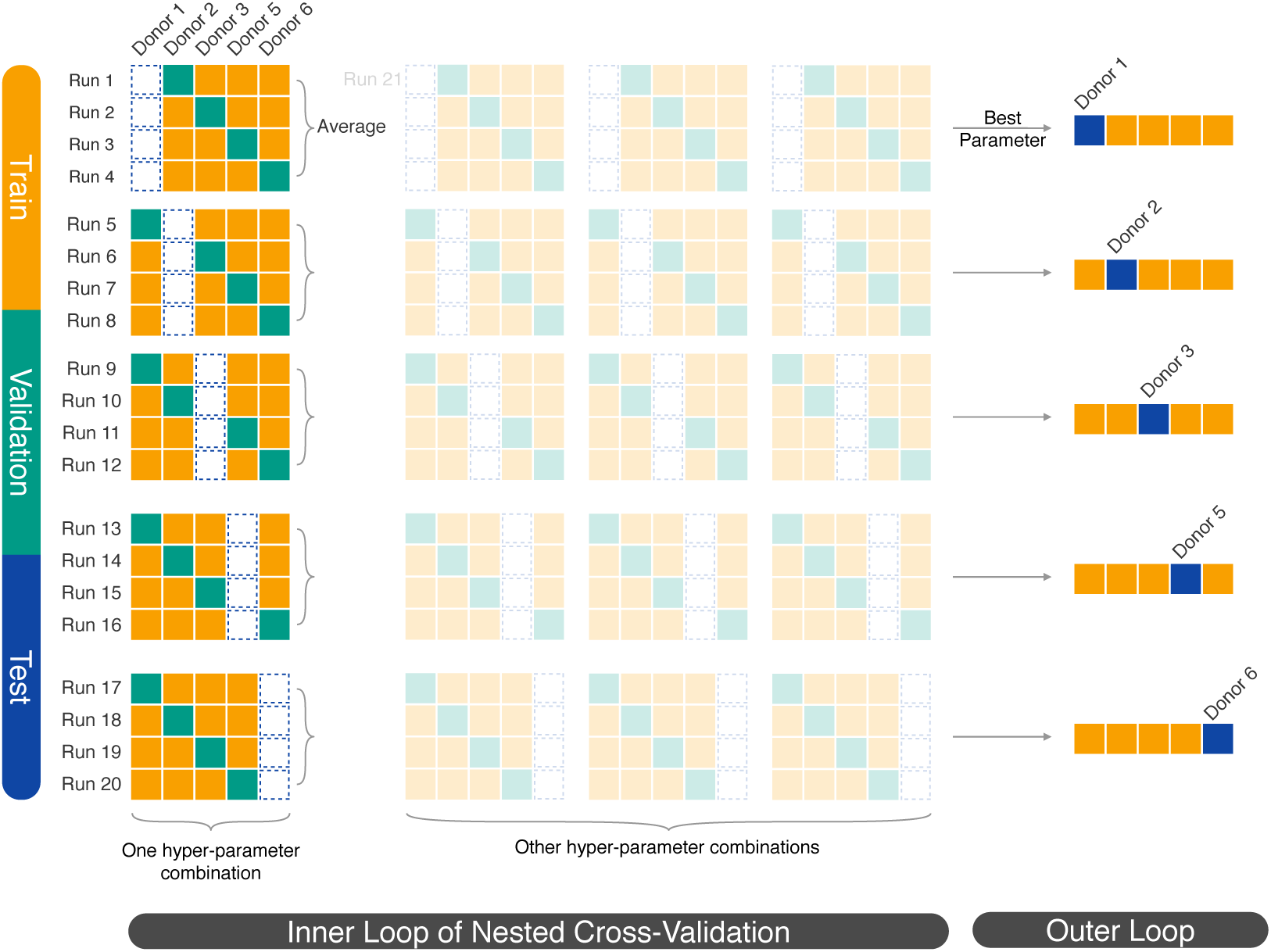
5 × 4 nested cross-validation scheme. For each test donor (blue), we used an inner cross-validation loop to optimize the hyper-parameters. We trained a model for each hyper-parameter combination using the training donors’ augmented images (yellow) and selected the hyper-parameters that performed best on the validation donor’s images (green). The validation donor is sometimes referred to as a tuning donor in cross-validation. Then, we trained a final model for each test donor using the selected hyper-parameters.

No single evaluation metric can capture all the strengths and weaknesses of a classifier, especially because our dataset was class imbalanced and not skewed in the same way for all donors. Therefore, we considered multiple evaluation metrics in the outer loop. Accuracy measures the percentage of correct predictions. It is easy to interpret, but it does not necessarily characterize a useful classifier. For example, when positive samples are rare, a trivial classifier that predicts all samples as negative yields high accuracy. Precision and recall (sensitivity), on the other hand, consider the costs of false positive and false negative predictions, respectively. Graphical metrics like the ROC curve and PR curve avoid setting a specific classification threshold. We used AUC to summarize ROC curves and average precision for the PR curves. The ROC curve is independent of the class distribution, while the PR curve is useful when the classes are imbalanced^35^. For this reason, we used mean average precision of the inner loop 4-fold cross-validation to select optimal hyper-parameters.

During the nested cross-validation, we trained the LeNet CNN and pre-trained CNN with fine-tuning using GPUs. These jobs ran on GTX 1080, GTX 1080 Ti, K40, K80, P100, or RTX 2080 Ti GPUs. All other models were trained using CPUs.

### Linear Classifiers

We used a trivial frequency classifier as a baseline model. This model computes the positive sample percentage in the training set. Then, it uses this frequency as a positive class prediction score (between 0 and 1) for all samples in the test set.

Logistic regression with Lasso regularization is a standard and interpretable statistical model used to classify microscopy images^8^. The Lasso approach reduces the number of effective parameters by shrinking the parameters of less predictive features to zero. These features are ignored when making a new prediction. We fitted and tested three Lasso logistic regression models with different types of features using the Python package scikit-learn^53^. An image intensity matrix with dimension 82 × 82 and values from 0 to 255, reshaped into a vector with length 6724, was used to fit the first model. The second model was trained with two scalar features, cell size and image total intensity, where cell size was computed using the pixel count in the cell mask generated by CellProfiler. The last model used 123 features relating to cell intensity, texture, and area, which were extracted from cell images using a CellProfiler pipeline with modules *MeaureObjectSizeShape, MeasureObjectIntensity*, and *MeasureTexture*. The Lasso regularization parameter *λ* was tuned for all three classifiers with nested cross-validation (Table S9). We also applied inverse class frequencies in the training data as class weights to adjust the imbalanced dataset.

### Simple Neural Network Classifiers

We developed a simple fully-connected neural network with one hidden layer (Fig. S16) using the Python package Keras with the TensorFlow backend^54, 55^. The input layer uses the flattened image pixel vector with dimension 6724 × 1. Network hyper-parameters – hidden neuron numbers, learning rate, and batch size – were tuned using nested cross-validation (Table S9). The cross-entropy loss function was weighted according to the class distribution in the training set.

Also, we trained a CNN with the LeNet architecture^33^ with randomly initialized weights (no pre-training). The LeNet architecture has two convolutional layers and two pooling layers (Fig. S17). We used the default number of neurons specified in the original paper in each layer. The input layer was modified to support 82 × 82 one-channel images, so we could train this network with image pixel intensities. Similar to the fully-connected neural network, we used nested cross-validation to tune the learning rate and batch size (Table S9) and applied class weighting. We used early stopping with a patience of 10 for both models, which means we stopped training if the loss function failed to improve on the early stopping set in 10 consecutive epochs.

### Pre-trained CNN Classifiers

We developed a transfer learning classifier that uses the Inception v3 CNN with pre-trained ImageNet weights^56, 57^. Instead of training the whole network end-to-end from scratch, we took advantage of the pre-trained weights by extracting and modeling off-the-shelf features or fine-tuning the last *n* Inception modules, where *n* was treated as a hyper-parameter (Fig. 9). Inception modules are mini-networks that constitute the overall Inception v3 architecture. The first approach is a popular practice for transfer learning with Inception v3. We freeze the weights of all layers before the output layer and use them to extract generic image characteristics. Then, we train a light-weight classifier from scratch, specifically a neural network with an average pooling layer and a fully connected hidden layer with 1024 neurons, using these off-the-shelf features. We refer to this model as the pre-trained CNN off-the-shelf model. An alternative is to fix some earlier layers and fine-tune the higher-level *n* layers by initializing them with pre-trained weights and continuing training on a new dataset. For this model, we modified the output layer to support binary classification, and we did not add new layers. In addition, we used the nested cross-validation scheme to optimize *n* along with the learning rate and batch size (Table S9), creating the pre-trained CNN with fine-tuning.

**Figure 9.**
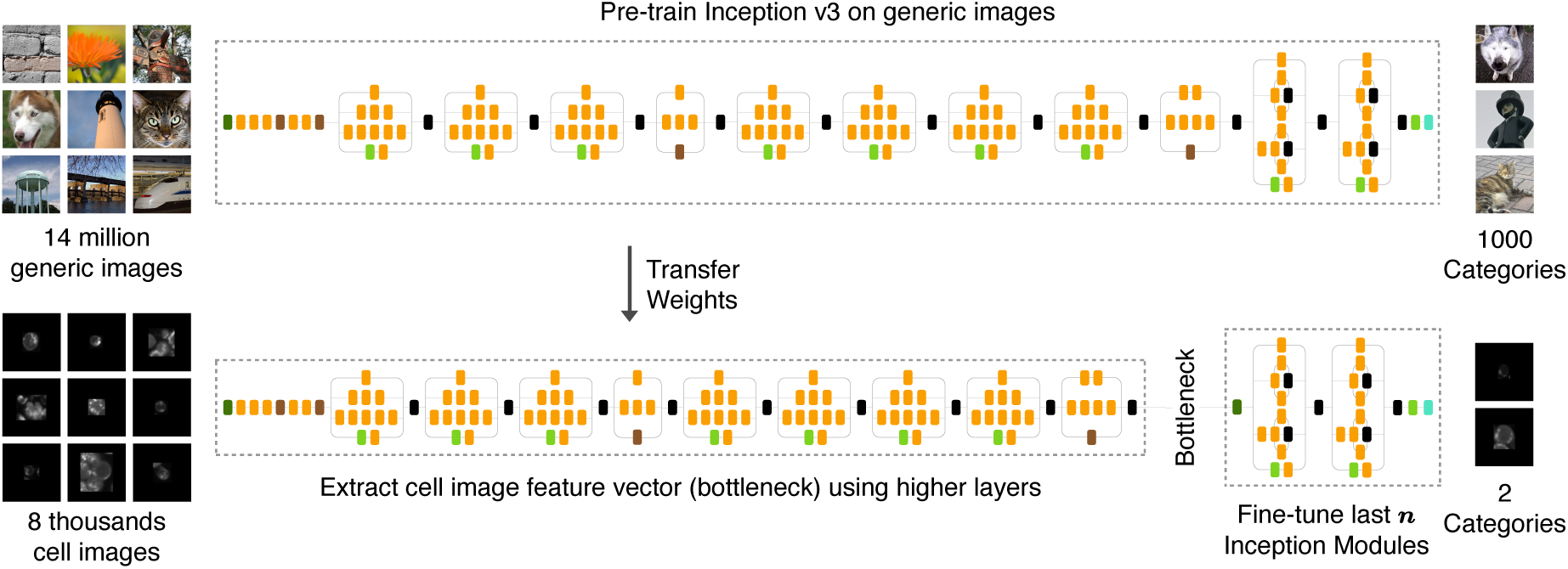
Fine-tuning the Inception v3 CNN to predict T cell activity. The example generic images are adapted from ImageNet.

To implement these two pre-trained CNN models, we resized the padded cell images with bilinear interpolation to fit the input layer dimension (299 × 299 × 3) and generated three-channel images by merging three copies of the same grayscale image. For the pre-trained CNN with fine-tuning, we first used the resized cell images to generate intermediate features (“bottlenecks”). Then, we used these features to fine-tune a sub-network. This approach significantly shortened the training time. Finally, we used class weighting and early stopping with a patience of 10 for both models. We implemented these two models using Keras with the TensorFlow backend.

### Pre-trained CNN Interpretation

We implemented multiple approaches for interpreting the pre-trained CNNs. Computing classification confidence on misclassified images can help us understand why classifiers make certain errors. The Softmax score is sometimes used as a confidence prediction. Softmax is a function that maps the output real-valued number (Logit) from a neural network into a score between 0 and 1, which is then used to make a classification as a class probability. However, using the Softmax score from a neural network as a confidence calibration does not match the real accuracy^36^. Therefore, we used temperature scaling to better calibrate the predictions^36^. After training, for each donor, we optimized the temperature *T* on the nested cross-validation outer loop validation set. Then, we applied *T* to scale the Logit before Softmax computation and used the new Softmax score to infer classification confidence.

In addition to confidence calibration, we used dimension reduction to investigate the high-dimensional representations learned by our pre-trained CNN models. Dimension reduction is a method to project high-dimensional features into lower dimensions while preserving the characteristics of the data. Therefore, it provides a good way to visualize how trained models represent different cell image inputs. In our study, we choose UMAP^37, 58^ as our dimension reduction algorithm. UMAP uses manifold learning techniques to reduce feature dimensions. It arguably preserves more of the global structure and is more scalable than the standard form of t-SNE^38^, an alternative approach. Using UMAP, we projected the image features, extracted from the CNN layer right before the output layer, from 2048 dimensions to two dimensions. We used the default UMAP parameter values: “n neighbors” as 15, “metrics” as “euclidean”, and “min dist” as 0.1. Then, we visualized and analyzed these projected features of T cell images using 2D scatter plots. When comparing UMAP with t-SNE, we used the default t-SNE parameters: “perplexity” as 30 and “metric” as “euclidean”.

For the pre-trained CNN with fine-tuning, each test donor has different tuned hyper-parameters and a different fine-tuned CNN. Therefore, we performed feature extraction and dimension reduction independently for each test donor. There is no guarantee that these five scatter plots share the same 2D basis. In contrast, the image pixel features, CellProfiler features, and off-the-shelf last layer features from a pre-trained CNN do not vary by test donor. For these three UMAP applications, we performed feature extraction and dimension reduction in one batch for all donors simultaneously. We excluded donor 4 from the dimension reduction analyses.

Finally, we used saliency maps to further analyze what morphology features were used in classification^39^. A saliency map is a straightforward and efficient way to detect how prediction value changes with respect to a small change in the input cell image pixels. It is generated by computing the gradient of the output class score with respect to the input image. We compared two ways to compute this gradient: standard backpropagation and guided backpropagation^40^. Backpropagation is a method to calculate the gradient of loss function with respect to the neural network’s weights. Guided backpropagation is a variant that only backpropagates positive gradients. We generated saliency maps of the output layer for the pre-trained CNN with fine-tuning model for test donor 1 with a few randomly sampled images from the test set. The saliency map interpretations help us assess whether the classification basis is intuitive and whether the predictions derive from image artifacts instead of cell morphology.

## Software and Data Availability

Our GitHub repository https://github.com/gitter-lab/t-cell-classification contains Jupyter note-books demonstrating how to run our Python code to pre-process images and train each of the classifiers. The notebooks can be run in a web browser using Binder. The software is available under the BSD 3-Clause Clear License. This repository also contains a randomly selected subset of the T cell images that can be used to quickly test our software. Our Zenodo dataset (DOI:10.5281/zenodo.2640835) contains bottleneck features from the Inception v3 model and trained model weights.

## Acknowledgements

We thank Tiffany Heaster for assistance with the T cell image processing; Quan Yin for CNN transfer learning advice; Shengchao Liu and Christine Walsh for general machine learning feedback; Katie Mueller, Steve Trier, and Kelsey Tweed for discussion of the classification results; and Jaime Frey and Zach Miller for assistance with the Cooley cluster. This research was funded by NIH R01 CA205101, the UW Carbone Cancer Center Support Grant NIH P30 CA014520, the Morgridge Institute for Research, and a UW-Madison L&S Honors Program Summer Senior Thesis Research Grant. In addition, this research benefited from GPU hardware from NVIDIA, resources of the Argonne Leadership Computing Facility, which is a DOE Office of Science User Facility supported under contract DE-AC02-06CH11357, the use of credits from the NIH Cloud Credits Model Pilot, a component of the NIH Big Data to Knowledge (BD2K) program, and the compute resources and assistance of the UW-Madison Center for High Throughput Computing (CHTC) in the Department of Computer Sciences. The CHTC is supported by UW-Madison, the Advanced Computing Initiative, the Wisconsin Alumni Research Foundation, the Wisconsin Institutes for Discovery, and the National Science Foundation and is an active member of the Open Science Grid.

## Supplemental Results

**Table S1.** Performance of Frequency Classifier.

| Donor | Accuracy | Precision | Recall | Average Precision | AUC | Activated Count | Quiescent Count |
| --- | --- | --- | --- | --- | --- | --- | --- |
| 1 | 13.16% | 13.16% | 100.00% | 13.16% | 50.00% | 235 | 1551 |
| 2 | 17.89% | 0.00% | 0.00% | 82.11% | 50.00% | 647 | 141 |
| 3 | 73.52% | 0.00% | 0.00% | 26.48% | 50.00% | 446 | 1238 |
| 5 | 26.48% | 0.00% | 0.00% | 73.52% | 50.00% | 683 | 246 |
| 6 | 56.75% | 0.00% | 0.00% | 43.25% | 50.00% | 442 | 580 |

**Table S2.**
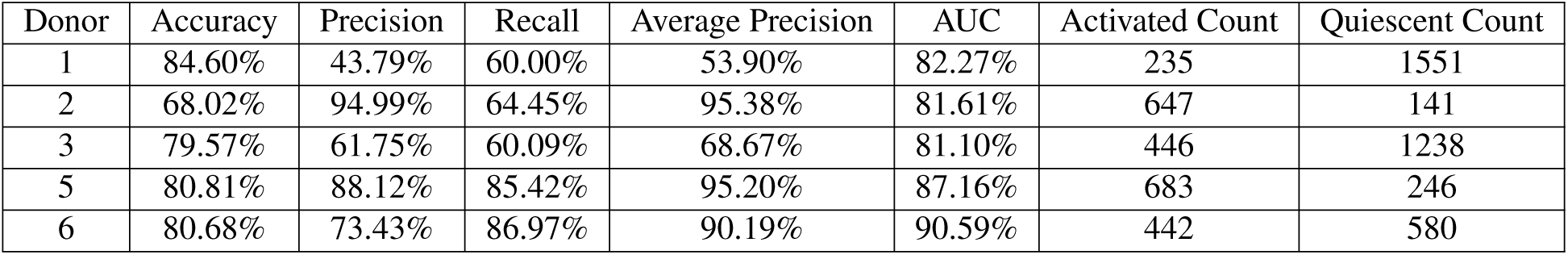
Performance of Logistic Regression (Image Pixel Matrix)

| Donor | Accuracy | Precision | Recall | Average Precision | AUC | Activated Count | Quiescent Count |
| --- | --- | --- | --- | --- | --- | --- | --- |
| 1 | 84.60% | 43.79% | 60.00% | 53.90% | 82.27% | 235 | 1551 |
| 2 | 68.02% | 94.99% | 64.45% | 95.38% | 81.61% | 647 | 141 |
| 3 | 79.57% | 61.75% | 60.09% | 68.67% | 81.10% | 446 | 1238 |
| 5 | 80.81% | 88.12% | 85.42% | 95.20% | 87.16% | 683 | 246 |
| 6 | 80.68% | 73.43% | 86.97% | 90.19% | 90.59% | 442 | 580 |

**Table S3.**
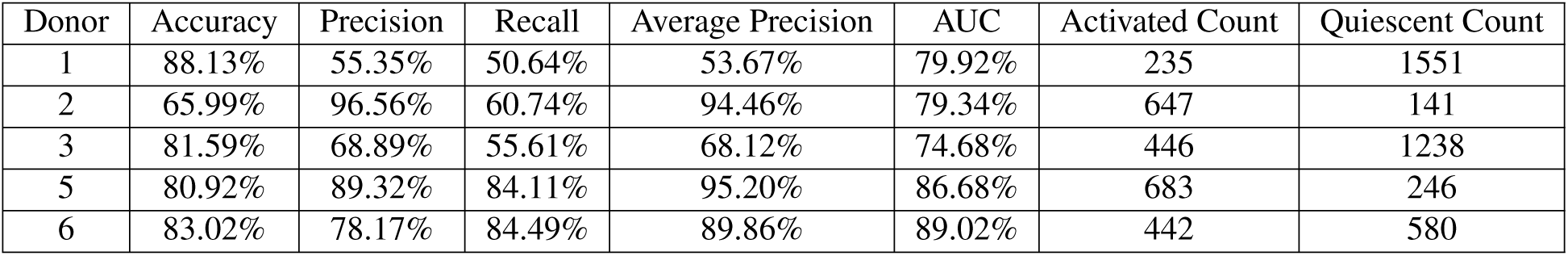
Performance of Logistic Regression (Total Intensity and Mask Size)

| Donor | Accuracy | Precision | Recall | Average Precision | AUC | Activated Count | Quiescent Count |
| --- | --- | --- | --- | --- | --- | --- | --- |
| 1 | 88.13% | 55.35% | 50.64% | 53.67% | 79.92% | 235 | 1551 |
| 2 | 65.99% | 96.56% | 60.74% | 94.46% | 79.34% | 647 | 141 |
| 3 | 81.59% | 68.89% | 55.61% | 68.12% | 74.68% | 446 | 1238 |
| 5 | 80.92% | 89.32% | 84.11% | 95.20% | 86.68% | 683 | 246 |
| 6 | 83.02% | 78.17% | 84.49% | 89.86% | 89.02% | 442 | 580 |

**Table S4.** Performance of Logistic Regression (CellProfiler Features)

| Donor | Accuracy | Precision | Recall | Average Precision | AUC | Activated Count | Quiescent Count |
| --- | --- | --- | --- | --- | --- | --- | --- |
| 1 | 95.74% | 86.98% | 79.57% | 88.85% | 95.61% | 235 | 1551 |
| 2 | 76.65% | 91.56% | 78.83% | 95.24% | 82.33% | 647 | 141 |
| 3 | 92.16% | 94.60% | 74.66% | 93.07% | 96.26% | 446 | 1238 |
| 5 | 81.81% | 81.81% | 96.78% | 93.74% | 86.70% | 683 | 246 |
| 6 | 89.33% | 82.33% | 95.93% | 95.97% | 97.01% | 442 | 580 |

**Table S5.** Performance of One-layer Fully Connected Neural Network.

| Donor | Accuracy | Precision | Recall | Average Precision | AUC | Activated Count | Quiescent Count |
| --- | --- | --- | --- | --- | --- | --- | --- |
| 1 | 88.80% | 55.28% | 75.74% | 65.40% | 90.55% | 235 | 1551 |
| 2 | 82.87% | 96.05% | 82.69% | 96.60% | 88.78% | 647 | 141 |
| 3 | 88.54% | 82.14% | 72.20% | 84.59% | 90.34% | 446 | 1238 |
| 5 | 84.78% | 85.58% | 95.19% | 96.48% | 92.05% | 683 | 246 |
| 6 | 87.41% | 80.62% | 93.48% | 94.44% | 95.36% | 442 | 580 |

**Table S6.** Performance of LeNet CNN.

| Donor | Accuracy | Precision | Recall | Average Precision | AUC | Activated Count | Quiescent Count |
| --- | --- | --- | --- | --- | --- | --- | --- |
| 1 | 94.23% | 78.21% | 77.87% | 82.15% | 95.36% | 235 | 1551 |
| 2 | 87.06% | 96.58% | 87.33% | 97.86% | 91.92% | 647 | 141 |
| 3 | 91.45% | 95.76% | 70.85% | 91.55% | 94.12% | 446 | 1238 |
| 5 | 87.41% | 87.83% | 96.19% | 96.40% | 92.41% | 683 | 246 |
| 6 | 87.38% | 78.61% | 97.29% | 96.70% | 97.36% | 442 | 580 |

**Table S7.**
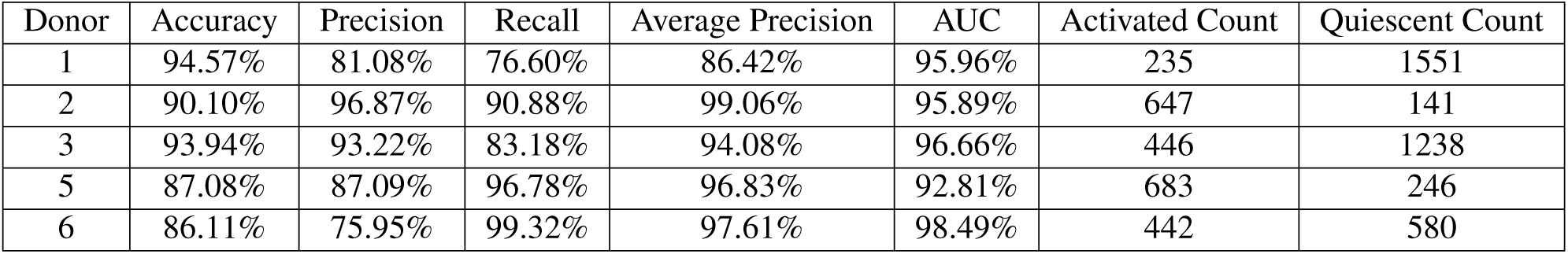
Performance of Pre-trained CNN Off-the-shelf Model.

| Donor | Accuracy | Precision | Recall | Average Precision | AUC | Activated Count | Quiescent Count |
| --- | --- | --- | --- | --- | --- | --- | --- |
| 1 | 94.57% | 81.08% | 76.60% | 86.42% | 95.96% | 235 | 1551 |
| 2 | 90.10% | 96.87% | 90.88% | 99.06% | 95.89% | 647 | 141 |
| 3 | 93.94% | 93.22% | 83.18% | 94.08% | 96.66% | 446 | 1238 |
| 5 | 87.08% | 87.09% | 96.78% | 96.83% | 92.81% | 683 | 246 |
| 6 | 86.11% | 75.95% | 99.32% | 97.61% | 98.49% | 442 | 580 |

**Table S8.** Performance of Pre-trained CNN with Fine-tuning.

| Donor | Accuracy | Precision | Recall | Average Precision | AUC | Activated Count | Quiescent Count |
| --- | --- | --- | --- | --- | --- | --- | --- |
| 1 | 96.81% | 92.79% | 82.13% | 91.71% | 95.70% | 235 | 1551 |
| 2 | 91.88% | 97.24% | 92.74% | 99.22% | 97.09% | 647 | 141 |
| 3 | 94.42% | 96.32% | 82.06% | 95.75% | 97.41% | 446 | 1238 |
| 5 | 89.77% | 89.20% | 97.95% | 97.02% | 94.26% | 683 | 246 |
| 6 | 94.91% | 92.76% | 95.70% | 98.20% | 98.91% | 442 | 580 |

**Table S9.**
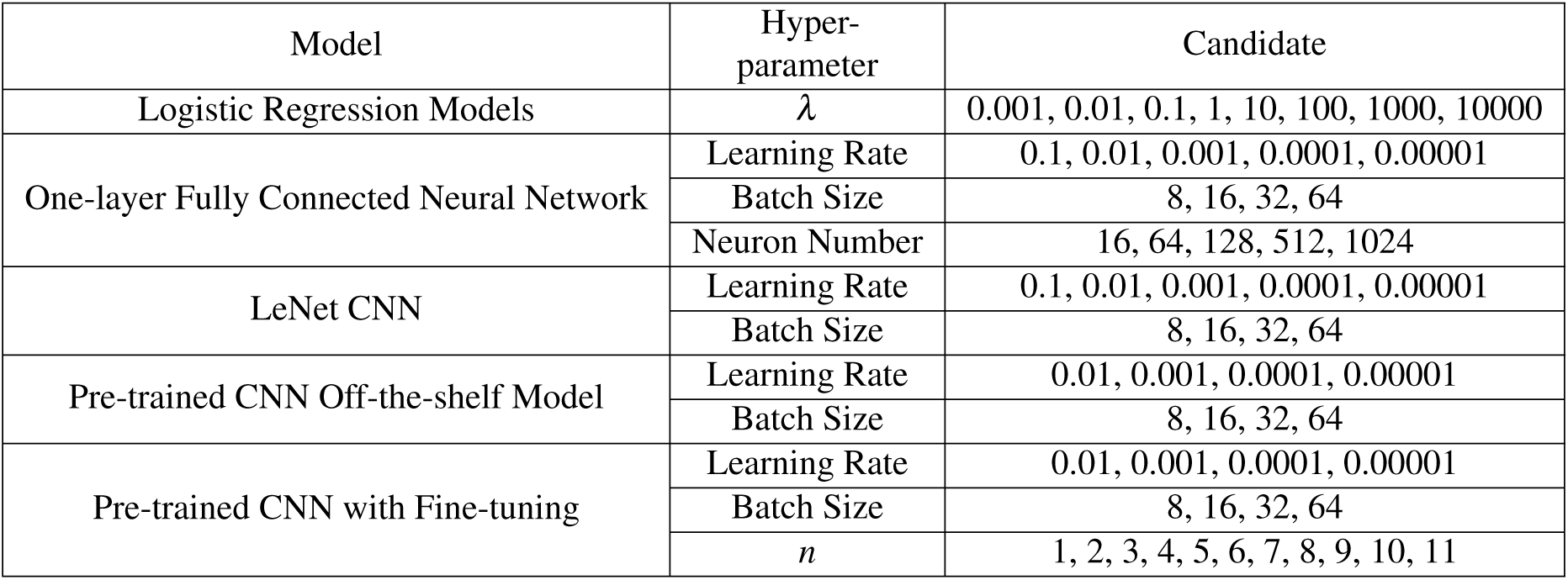
Parameter Candidates for Grid Search.

| Model | Hyper-parameter | Candidate |
| --- | --- | --- |
| Logistic Regression Models | $\lambda$ | 0.001, 0.01, 0.1, 1, 10, 100, 1000, 10000 |
| One-layer Fully Connected Neural Network | Learning Rate | 0.1, 0.01, 0.001, 0.0001, 0.00001 |
|  | Batch Size | 8, 16, 32, 64 |
|  | Neuron Number | 16, 64, 128, 512, 1024 |
| LeNet CNN | Learning Rate | 0.1, 0.01, 0.001, 0.0001, 0.00001 |
|  | Batch Size | 8, 16, 32, 64 |
| Pre-trained CNN Off-the-shelf Model | Learning Rate | 0.01, 0.001, 0.0001, 0.00001 |
|  | Batch Size | 8, 16, 32, 64 |
| Pre-trained CNN with Fine-tuning | Learning Rate | 0.01, 0.001, 0.0001, 0.00001 |
|  | Batch Size | 8, 16, 32, 64 |
| | $n$ | 1, 2, 3, 4, 5, 6, 7, 8, 9, 10, 11 |

**Figure S1.**
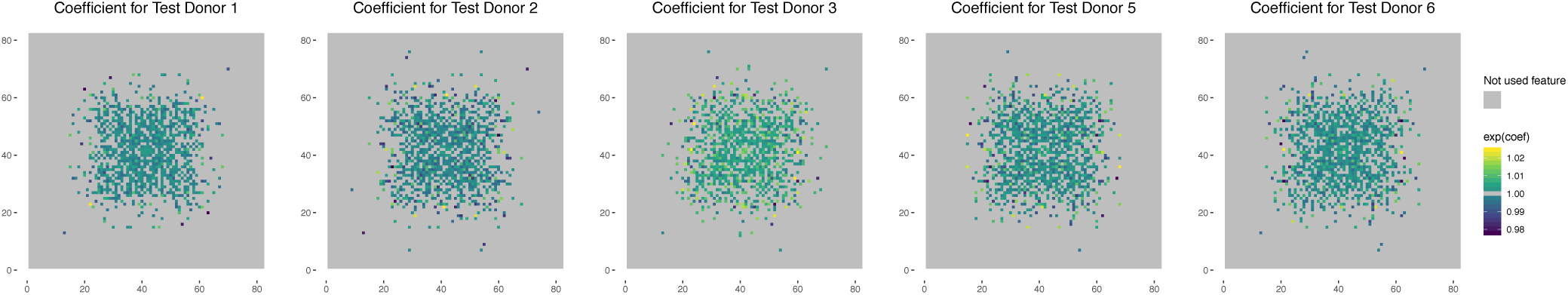
Visualization of odds ratios for one unit increase of each pixel’s intensity value. A pixel with odds ratio higher than 1 (light green to yellow) means that, fixing all other pixels, a one-unit intensity increase in this pixel leads to a higher likelihood of predicting the cell is active.

**Figure S2.**
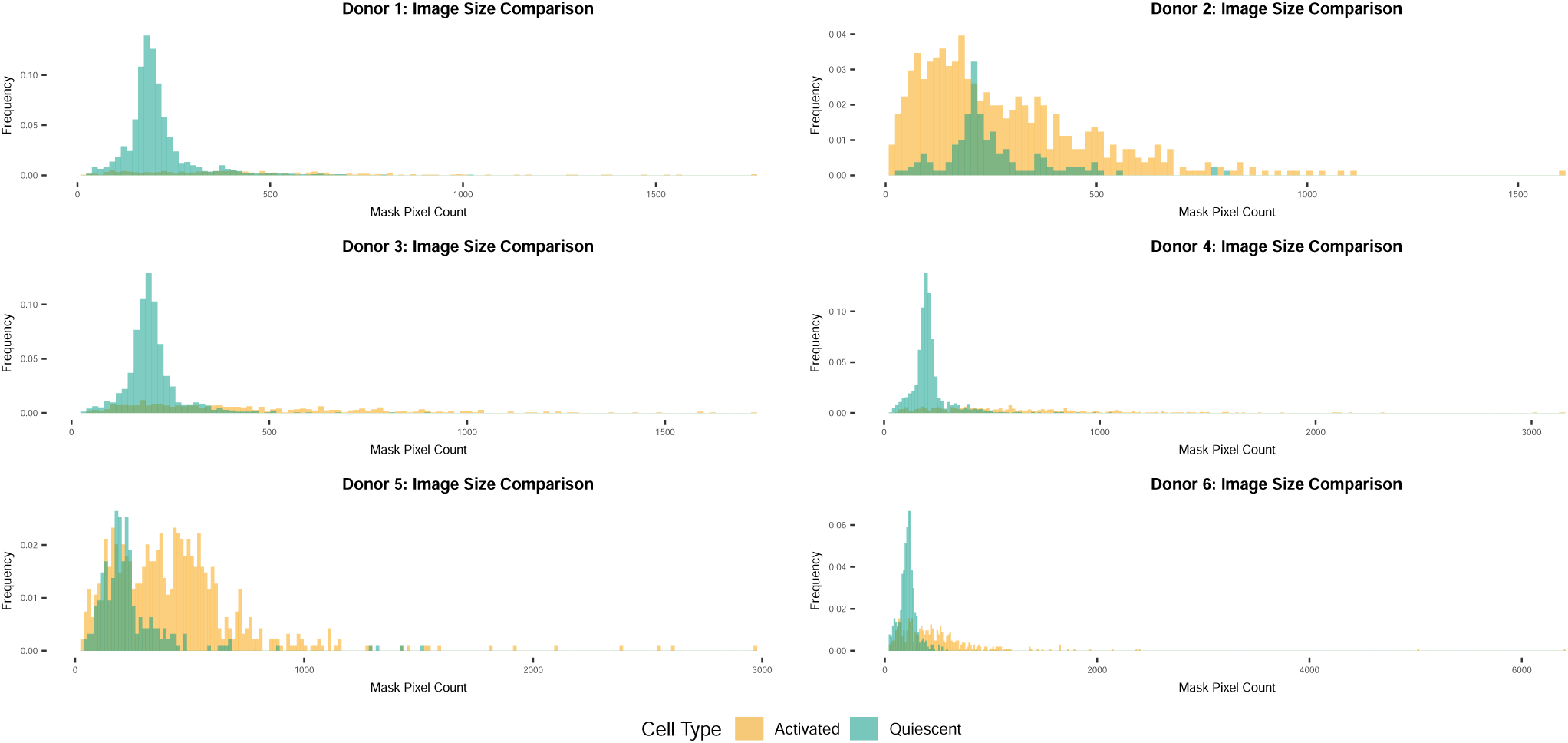
Mask size distribution for each of the donors.

**Figure S3.**
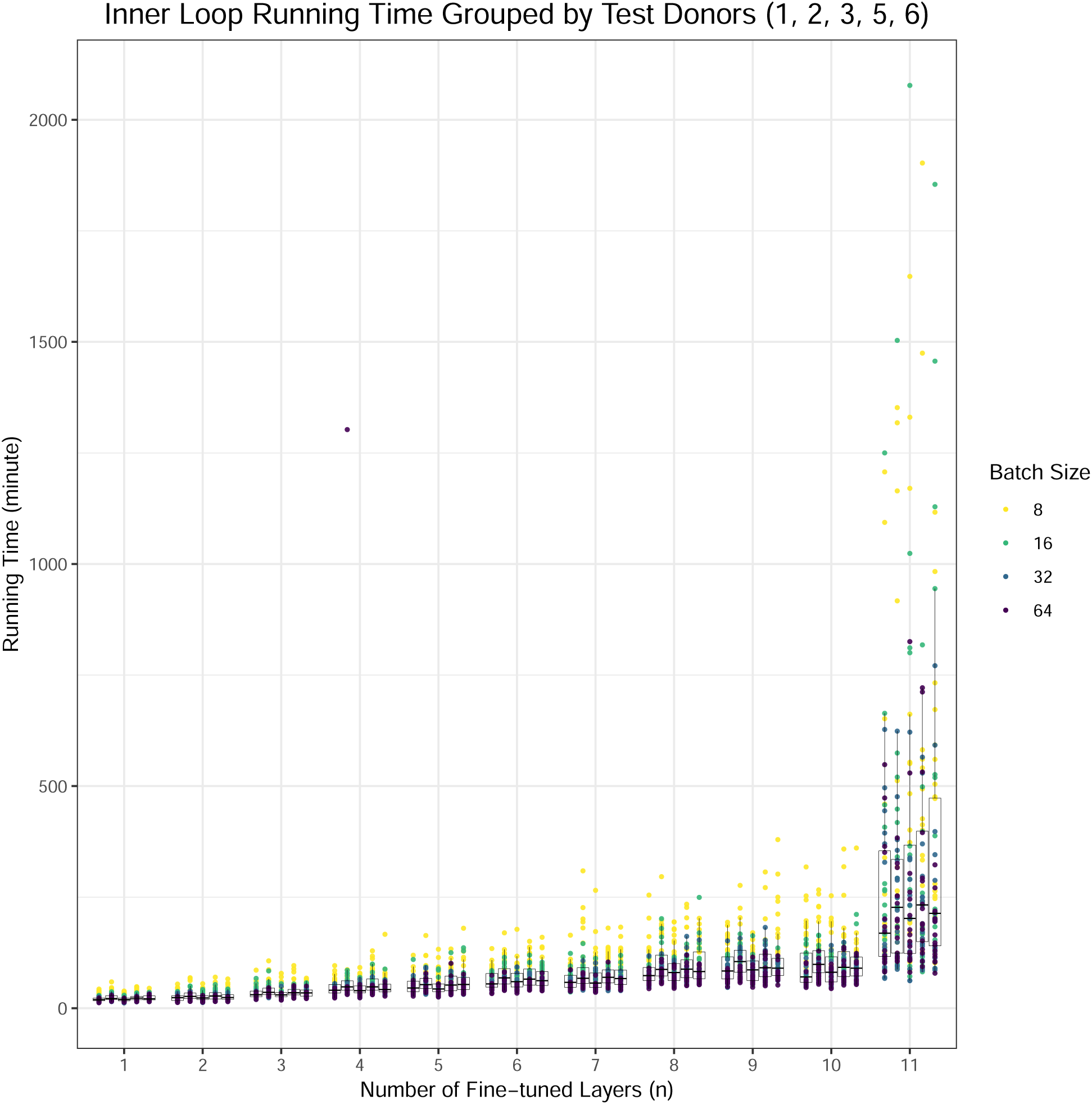
Nested cross-validation inner loop execution time for each *n*, test donor, and batch size.

**Figure S4.**
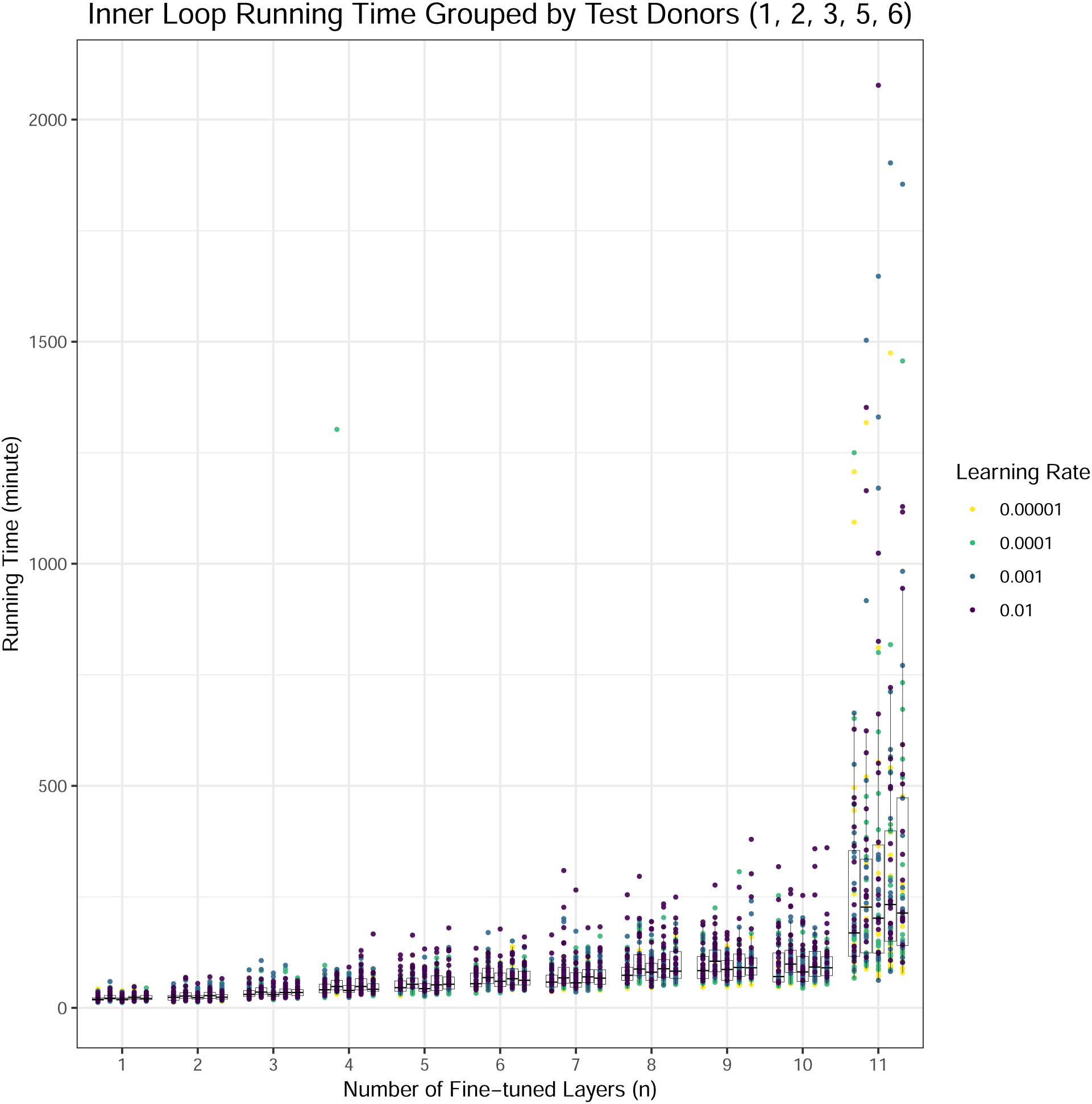
Nested cross-validation inner loop execution time for each *n*, test donor, and learning rate.

**Figure S5.**
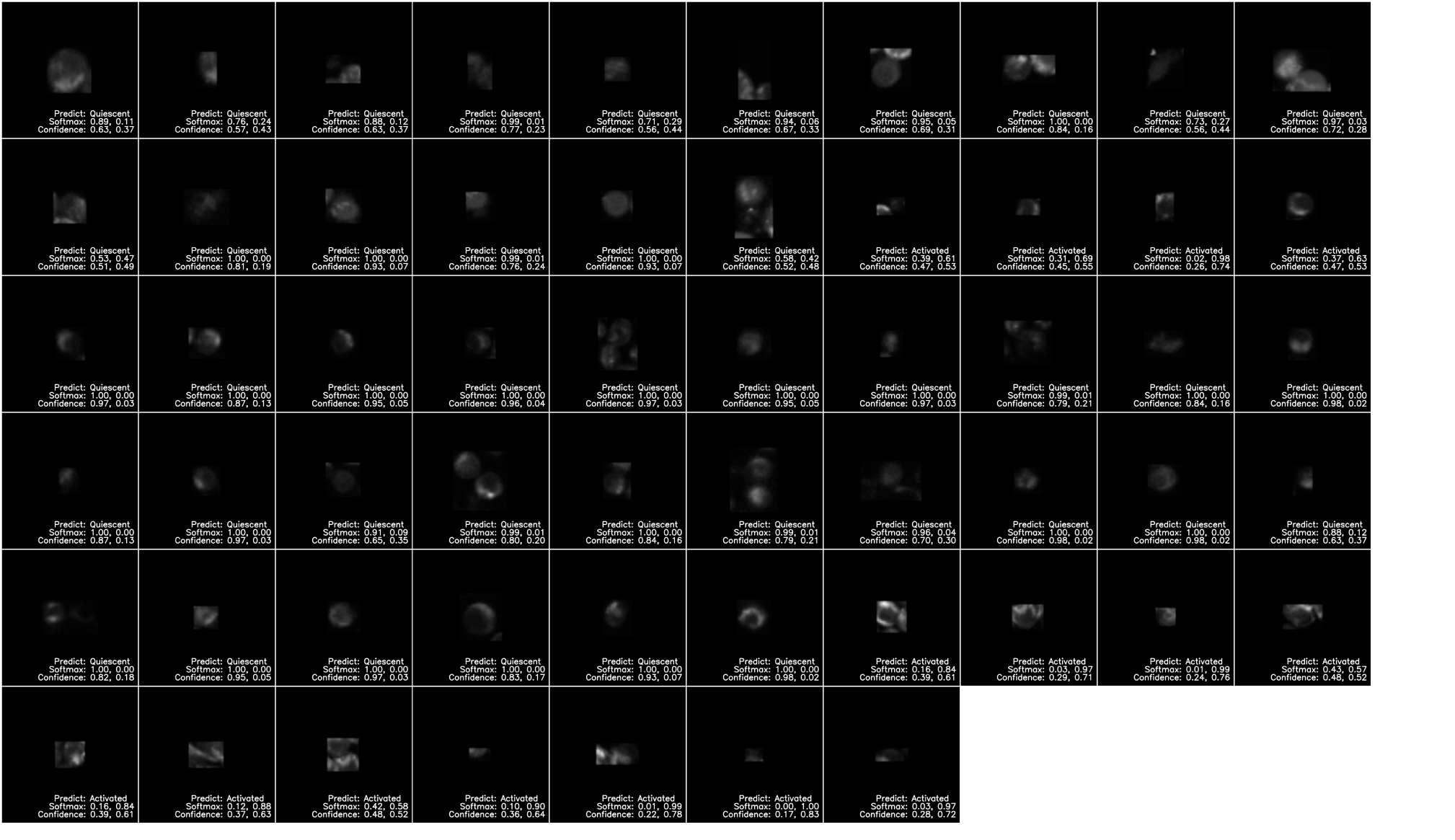
Pre-trained CNN with Fine-tuning Misclassified Images: Donor 1.

**Figure S6.**
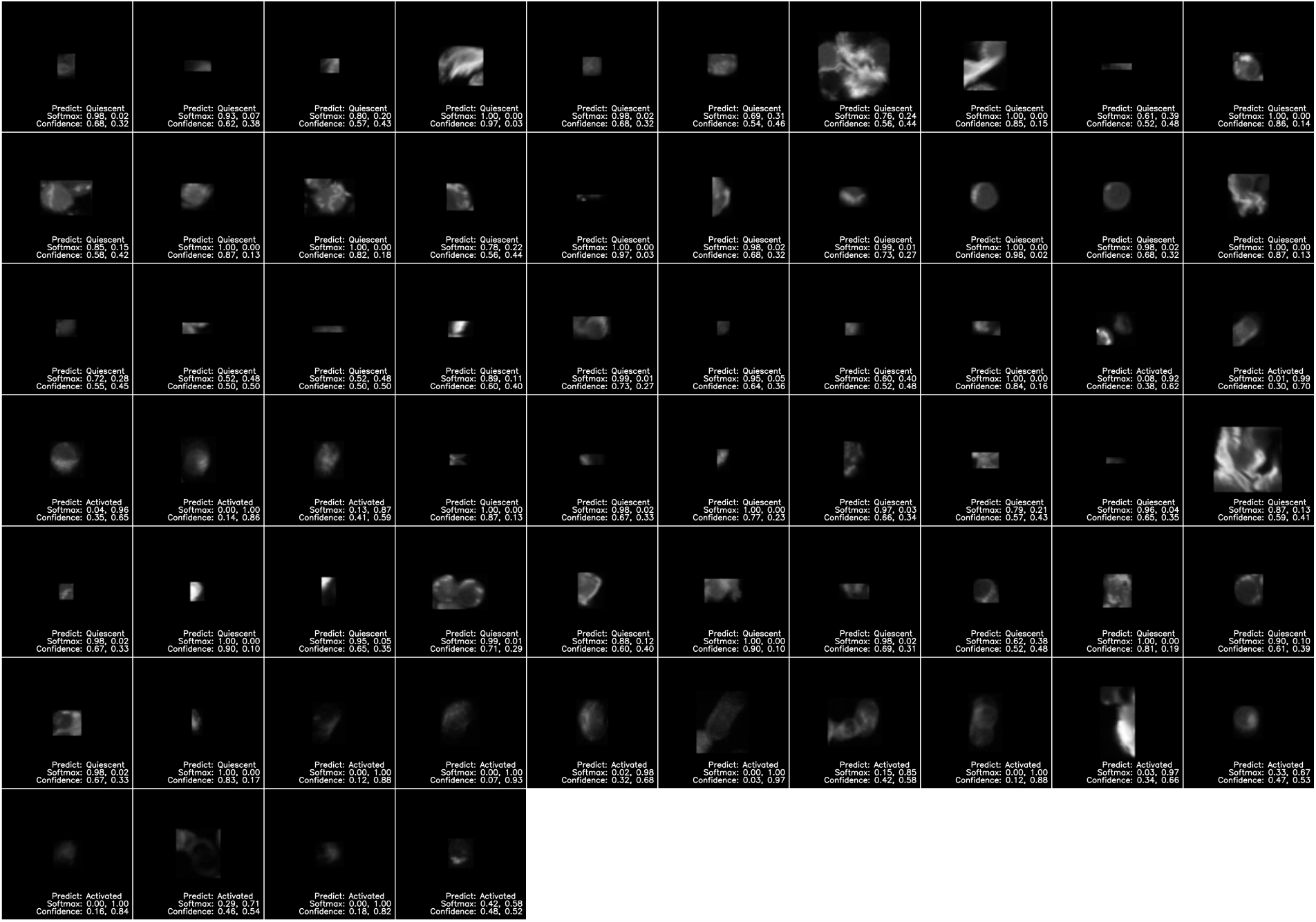
Pre-trained CNN with Fine-tuning Misclassified Images: Donor 2.

**Figure S7.**
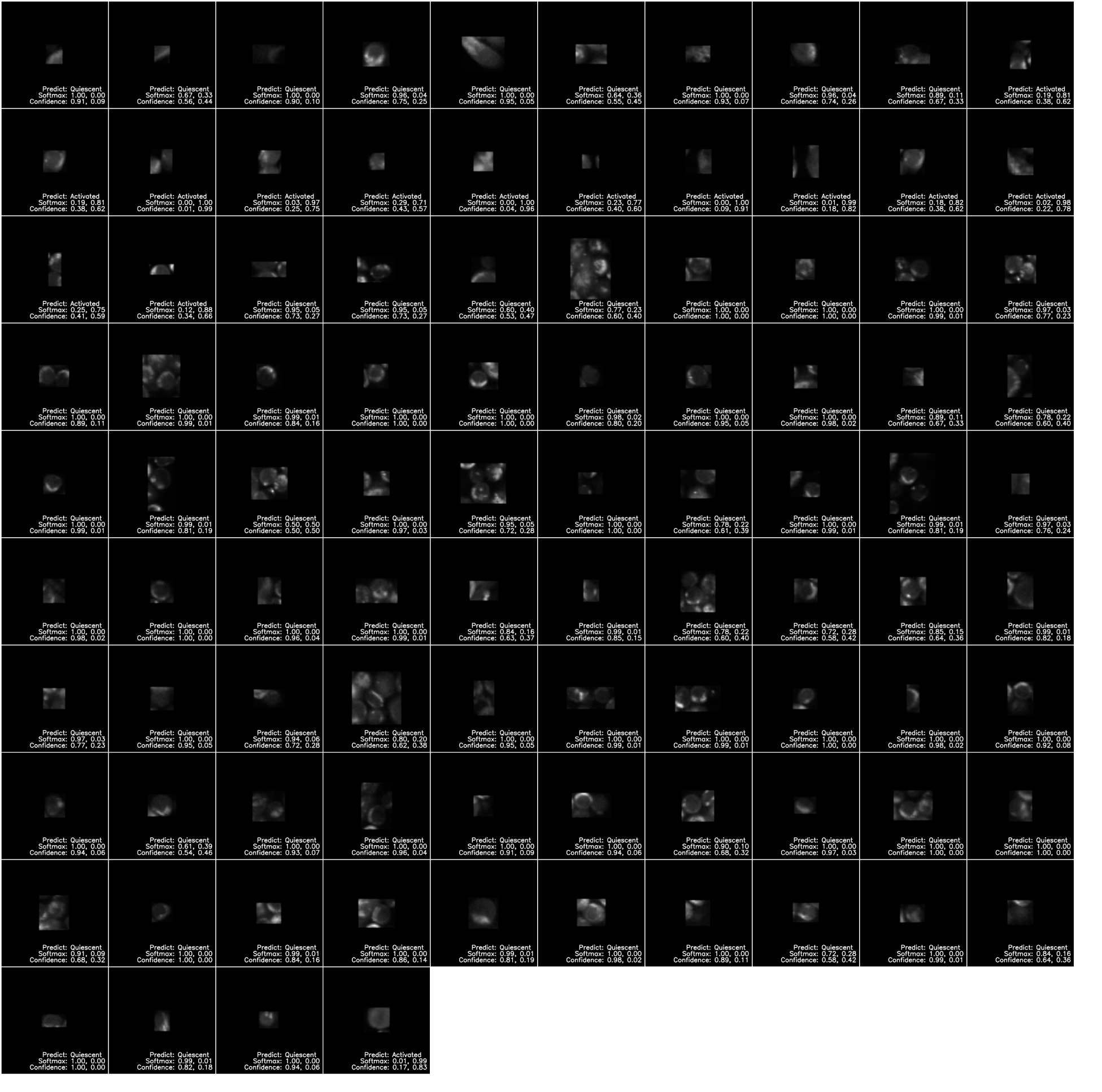
Pre-trained CNN with Fine-tuning Misclassified Images: Donor 3.

**Figure S8.**
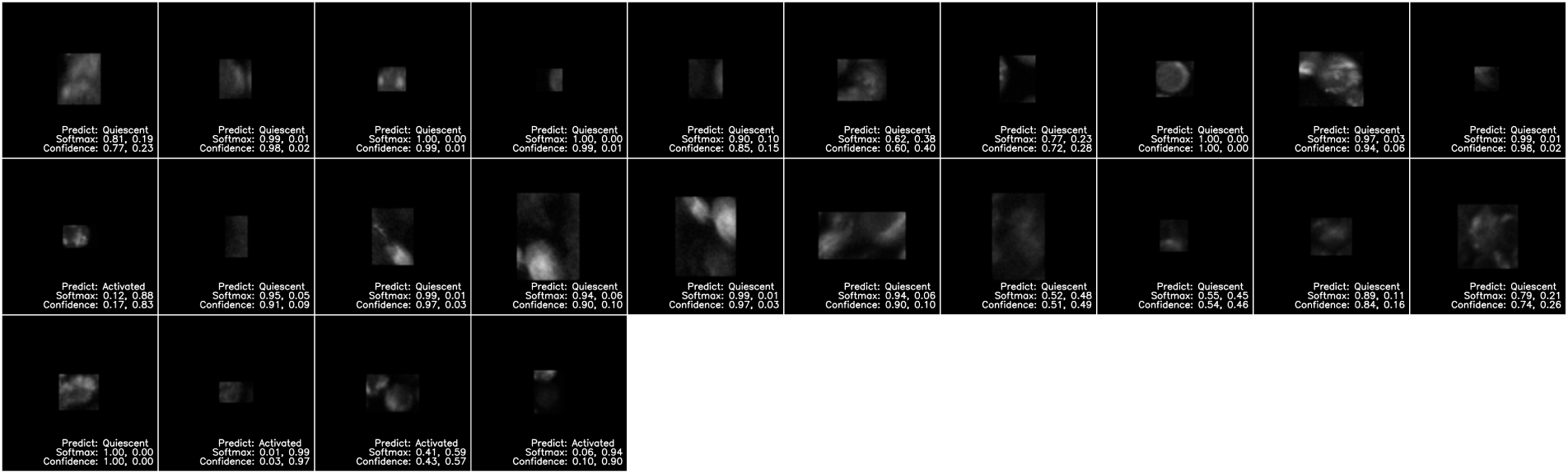
Pre-trained CNN with Fine-tuning Misclassified Images: Donor 4.

**Figure S9.**
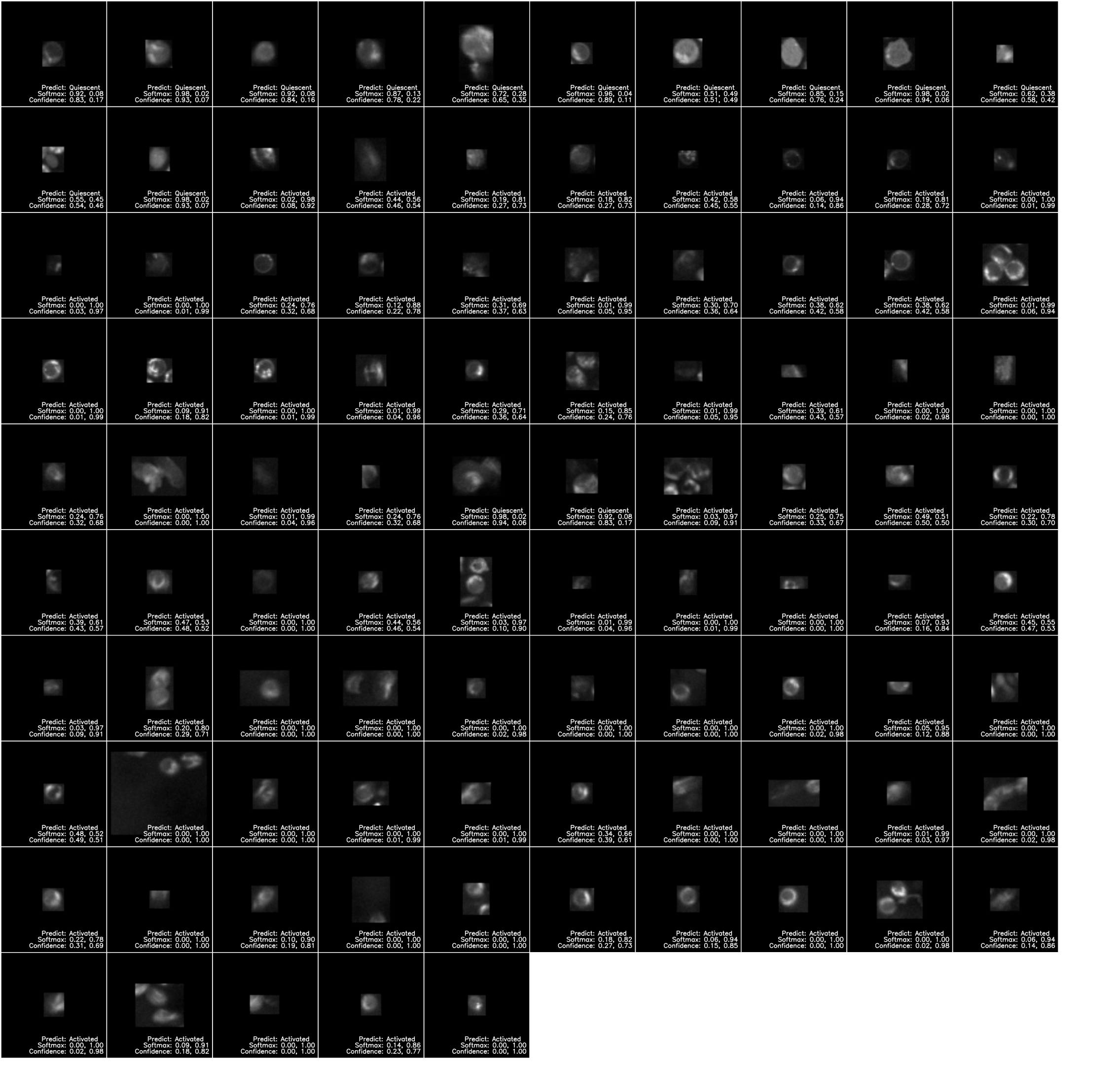
Pre-trained CNN with Fine-tuning Misclassified Images: Donor 5.

**Figure S10.**
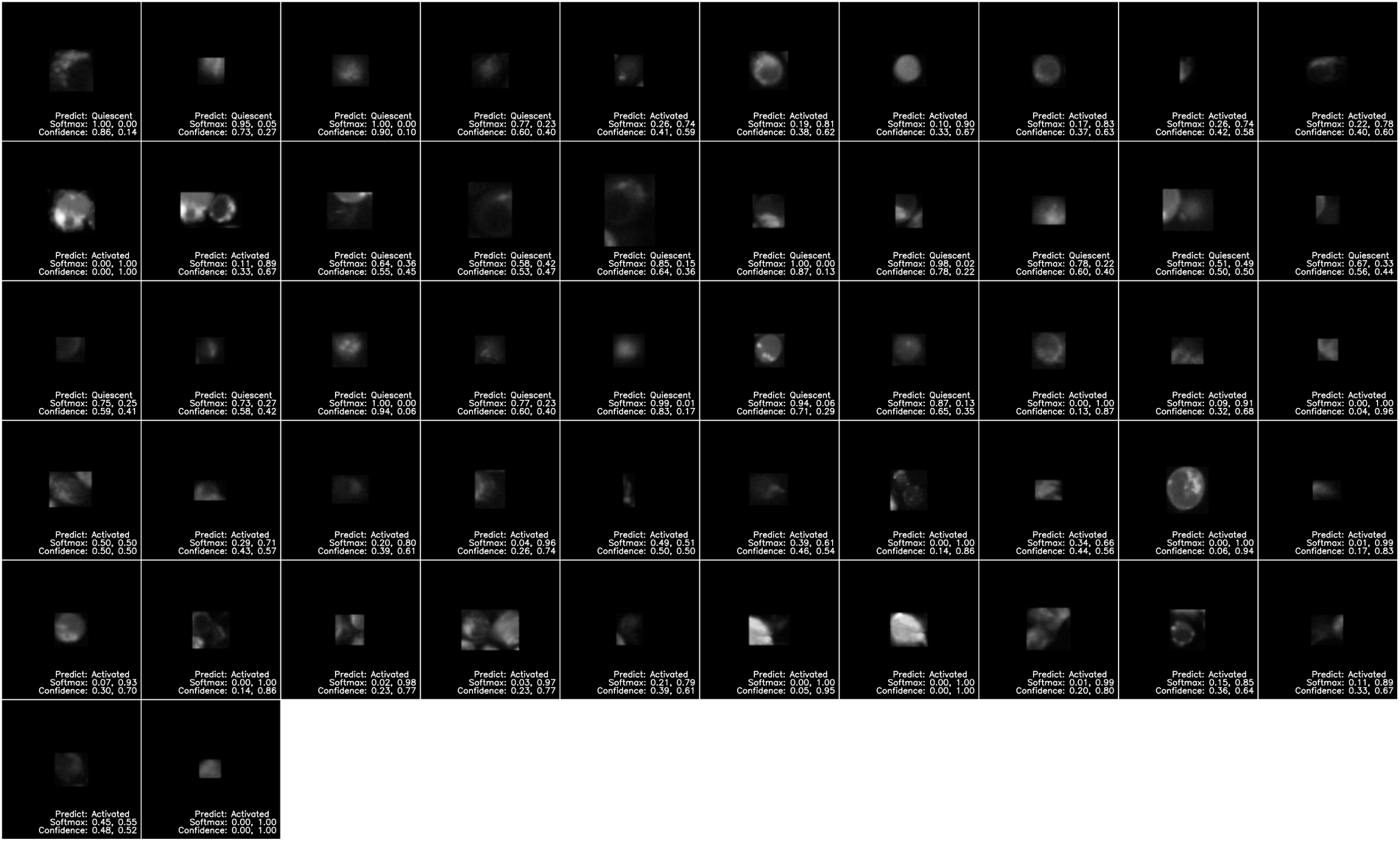
Pre-trained CNN with Fine-tuning Misclassified Images: Donor 6.

**Figure S11.**
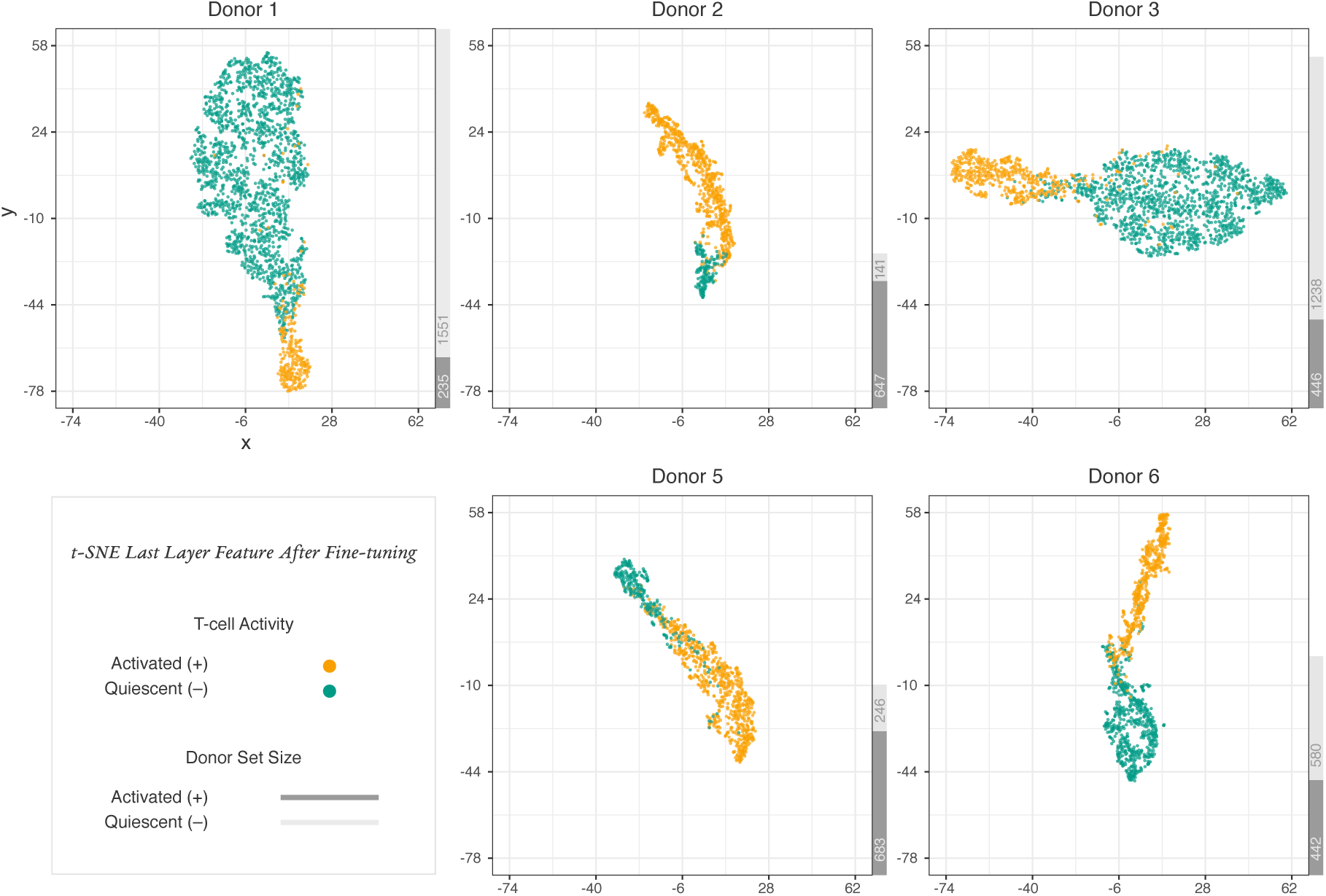
2D representations of T cell features extracted from the last layer of the pre-trained CNN with fine-tuning. Dimensions were reduced from 2048 using t-SNE.

**Figure S12.**
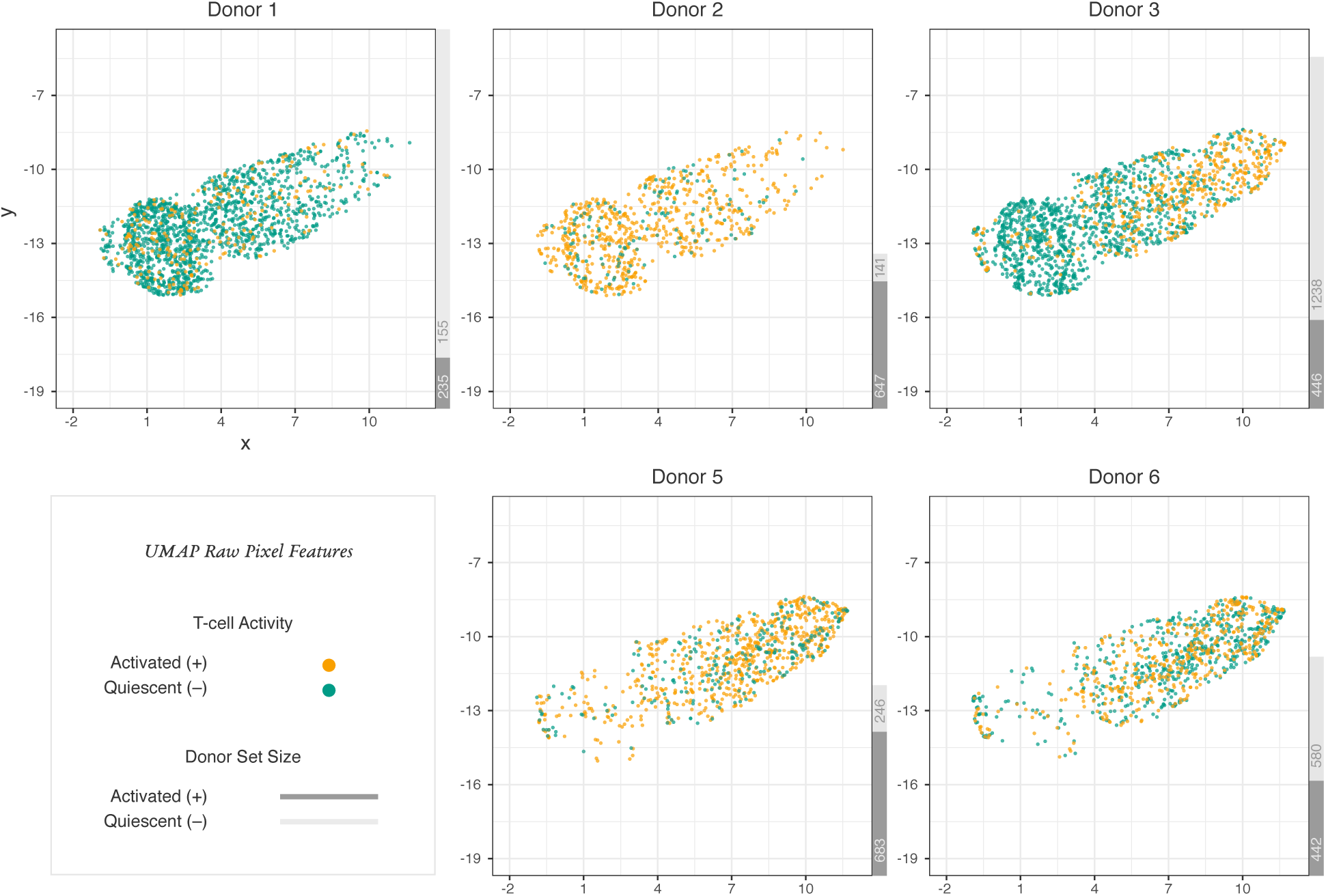
2D representations of T cell raw pixel features. Dimensions were reduced from 82 *×* 82 = 6724 using UMAP.

**Figure S13.**
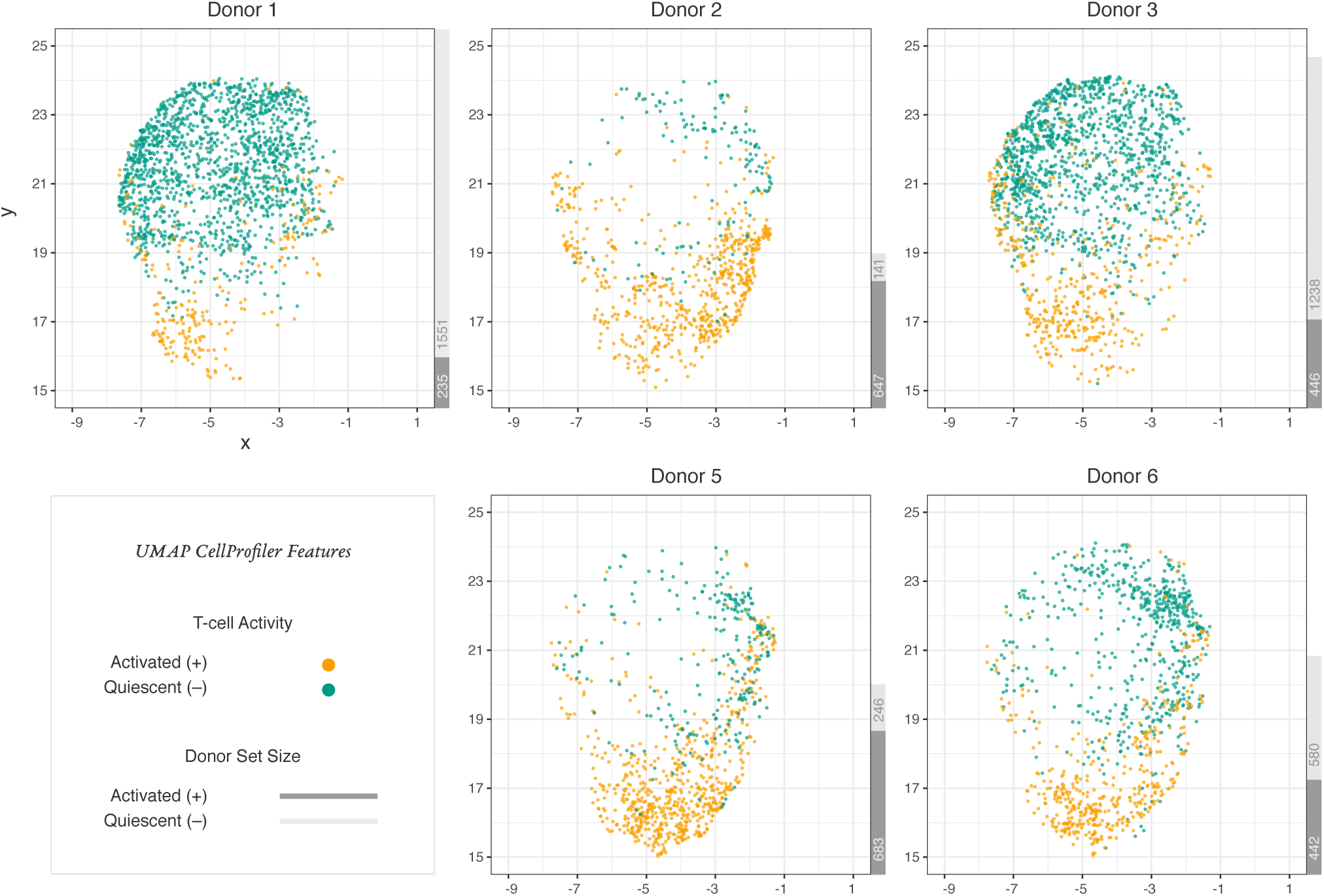
2D representations of T cell CellProfiler features. Dimensions were reduced from 123 using UMAP.

**Figure S14.**
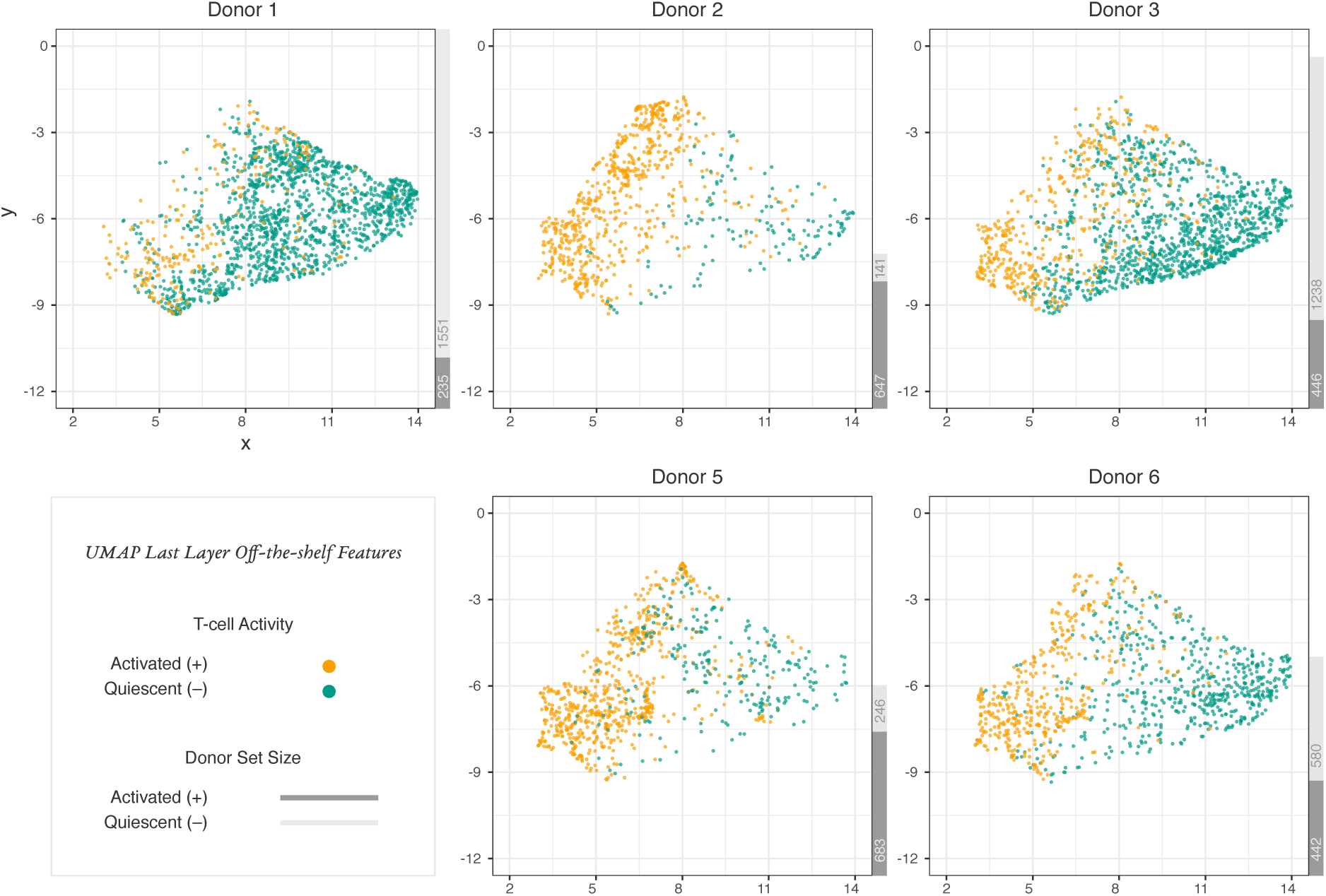
2D representations of T cell features extracted from the last layer of the pre-trained Inception v3 model before fine-tuning. Dimensions were reduced from 2048 using UMAP.

**Figure S15.**
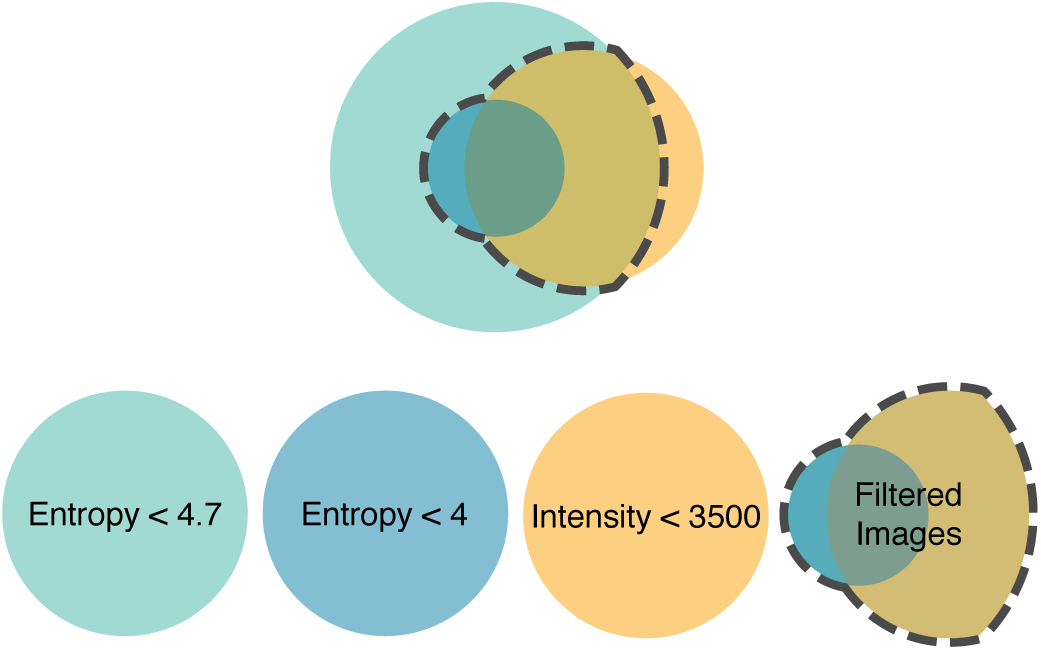
The segmented images are removed from the dataset if their entropy is less than 4 or if their entropy is less than 4.7 and their intensity is less than 3500.

**Figure S16.**
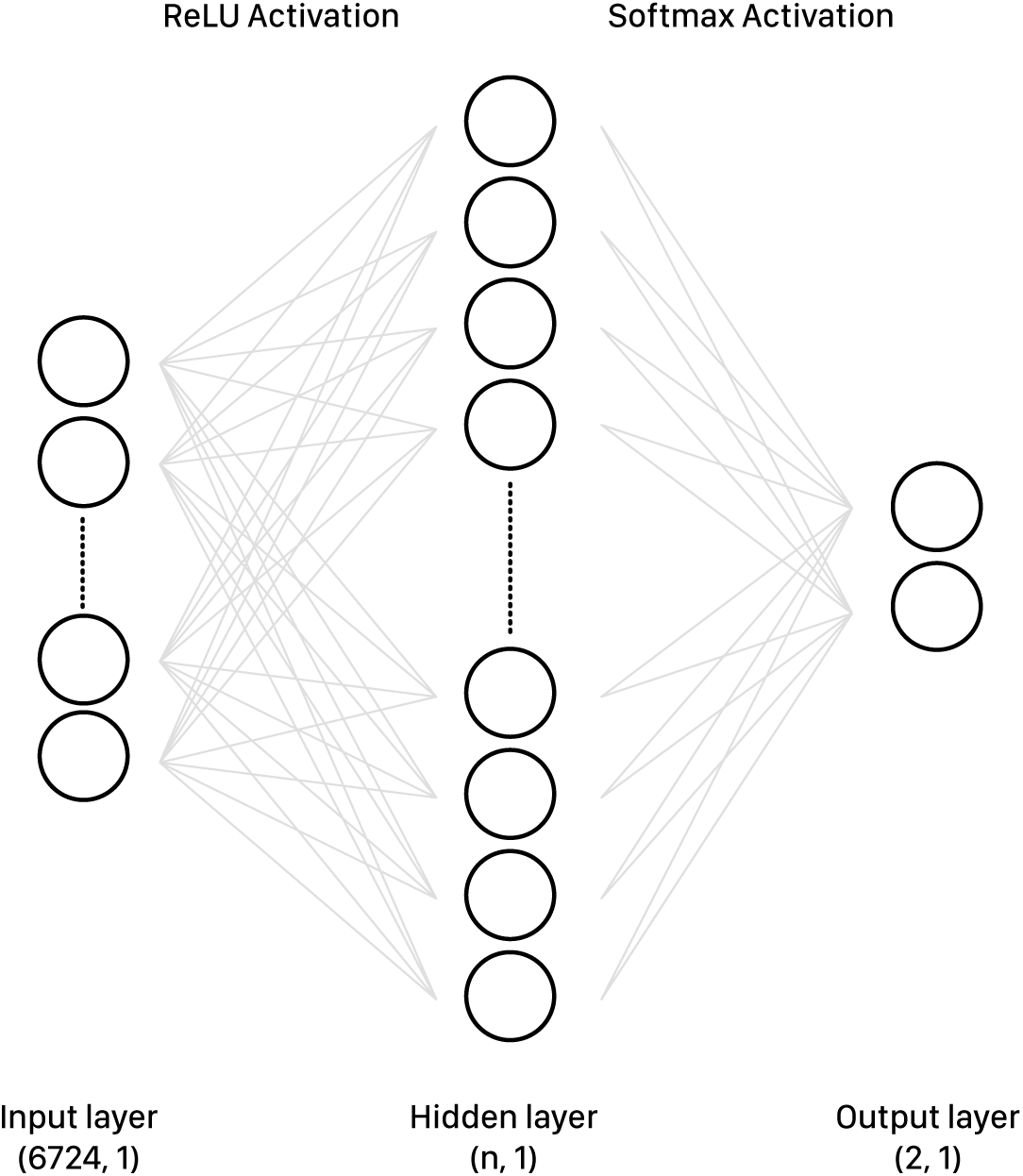
One-layer Fully Connected Neural Network architecture.

**Figure S17.**
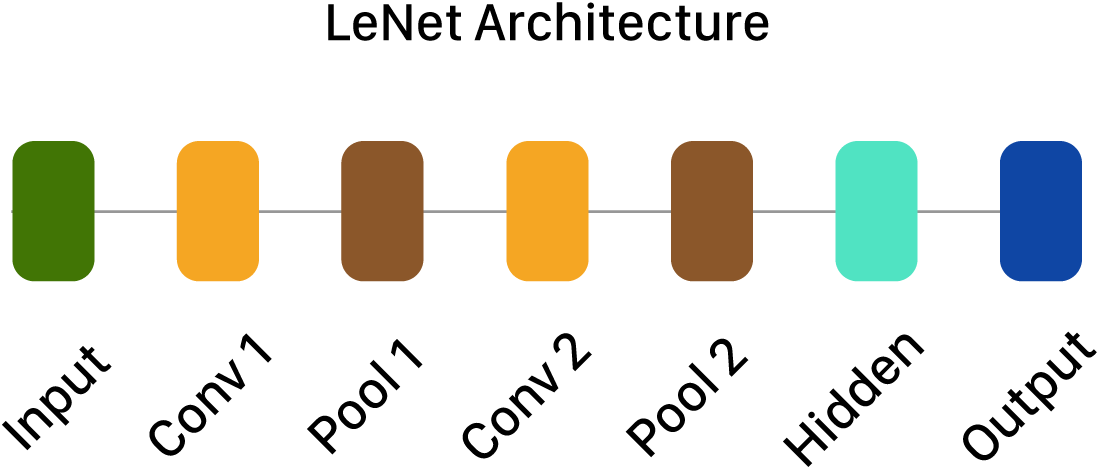
LeNet CNN architecture.

